# RNA-Stabilized Coat Proteins for Sensitive and Simultaneous Imaging of Distinct Single mRNAs in Live Cells

**DOI:** 10.1101/2024.11.21.624393

**Authors:** Christopher J. Kuffner, Alexander M. Marzilli, John T. Ngo

## Abstract

RNA localization and regulation are critical for cellular function, yet many live RNA imaging tools suffer from limited sensitivity due to background emissions from unbound probes. Here, we introduce conditionally stable variants of MS2 and PP7 coat proteins (which we name dMCP and dPCP) designed to decrease background in live-cell RNA imaging. Using a protein engineering approach that combines circular permutation and degron masking, we generated dMCP and dPCP variants that rapidly degrade except when bound to cognate RNA ligands. These enhancements enabled the sensitive visualization of single mRNA molecules undergoing differential regulation within various sub-compartments of live cells. We further demonstrate dual-color imaging with orthogonal MS2 and PP7 motifs, allowing simultaneous low-background visualization of distinct RNA species within the same cell. Overall, this work provides versatile, low-background probes for RNA imaging, which should have broad utility in the imaging and biotechnological utilization of MS2- and PP7-containing RNAs.

---

The regulation of messenger RNA (mRNA) abundance, localization, and translation is crucial for many cellular behaviors.^1,2^ These include ensuring correct targeting of proteins for secretion,^3^ controlling cell motility in response to external stimuli,^4^ and facilitating synaptic plasticity between connected neurons.^5^ Although many of these pathways rely on similar mechanisms of selective subcellular trafficking and protein translation, knowing where and when these processes occur is critical for understanding the causes and effects of mRNA regulation. As such, methods that allow researchers to visualize the localization and dynamics of individual, single mRNA transcripts have become valuable tools for investigating RNA biology.

User-friendly imaging probes that can be readily implemented in living specimens are especially important tools in resolving the dynamics of mRNAs in real time.^6^ While a variety of such probes have been developed, those that are the most widely used are based on bacteriophage-derived components – primarily the MS2 and PP7 coat proteins (MCP and PCP) and their cognate MS2 and PP7 RNA hairpins.^7–9^ By expressing fluorescent protein (FP)-fused MCP or PCP units, researchers can track single transcripts tagged with MS2 and/or PP7 arrays.

However, selectively visualizing tagged RNAs with these probes can be challenging due to background emissions from unbound coat protein copies, which can hinder the detection of RNA-bound species. To overcome this challenge, researchers have exploited nuclear localization signals (NLSs) to direct and sequester unbound coat protein units within the nucleus, allowing mature mRNAs to be visualized with increased contrast in the cytoplasm.^7^ Alternatively, RNA-specific contrast can be enhanced by increasing the number of fluorophores targeted to a given mRNA sequence.^10–12^. While these strategies can enhance the detectability of tagged RNAs, they do not offer a solution to the broader challenge of mismatched coat protein-stem loop stoichiometries---a primary limiting factor in the specificity and utility of MS2- and PP7-based technologies.

Recently, RNA-responsive reporter systems, such as fluorogenic RNA aptamers^13–15^ and RNA-templated protein complementation^16–18^, have been developed as alternatives to traditional coat protein-based strategies. In these approaches, signal generation is made RNA-dependent, thus allowing users to track tagged RNAs in detail and quantify their levels throughout cells. However, practical limitations make implementing these probes challenging in certain experimental contexts. For example, aptamer-based systems require exogenous chromophores, which can be challenging to supply and maintain in certain cells and model systems. These methods are further restricted due to obstacles associated with the limited brightness or the limited availability of their associated chromophores. These factors restrict the versatility of fluorogenic probes, especially compared to direct FP-coat protein fusions, in which FPs can be readily substituted with new sequences. Overall, an ideal system would combine the versatility of direct coat protein fusions with the reduced background and RNA-dependent signals of fluorogenic systems.

In more recent work, a labeling strategy that meets these criteria was developed based on so-called ‘fluorogenic proteins.’ In this approach, a conditionally stable RNA-binding protein is designed to be degraded unless bound to its target RNA, thereby reducing the levels of unbound proteins while rendering tagged RNA ‘fluorogenically’ visible via binding-induced reporter protein preservation.^19^ To develop such a probe, a virally-derived Tat peptide was modified to contain a C-terminal degron,^20^ resulting in a 19 amino acid sequence (called ‘tDeg’), which renders proteins unstable when fused to their C-termini while maintaining the ability to bind TAR-like RNA sequences (called ‘Pepper’). Critically, binding between tDeg-fused proteins and Pepper-tagged RNAs induces the selective stabilization of the RNA-bound form of tDeg-tagged proteins, an effect that is mediated via steric shielding of the degron by Pepper RNA, thus neutralizing its recognition by degradation machinery.

Recognizing the utility of multicolor RNA imaging, and motivated by a growing need for improved RNA-binding tools, we set out to complement the tDeg strategy by developing destabilized versions of MCP and PCP. In natural contexts, MCP and PCP bind their cognate hairpins via orientations in which their N- and C-terminal ends are solvent-exposed. Thus, to render them conditionally stable, we implemented a two-step approach in which (1) circular permutation was first used to reorient coat protein termini to RNA-adjacent locations, followed by (2) the attachment and positional optimization of RNA-maskable C-degrons (**Fig. 1a**). Using this approach, we generated a conditionally stable MCP that is efficiently degraded by cells while undergoing a more than 50-fold stability enhancement in response to MS2 binding. We confirmed the versatility of the domain by testing diverse fusions in fluorescence and bioluminescent assays. Furthermore, we exploited its RNA-dependent nature to sensitively visualize and record the dynamics of single mRNAs distributed throughout various subcellular locales, including within the nucleus and cytoplasm, on the surface of mitochondria, and as bound to the endoplasmic reticulum (ER).

**Figure 1.**
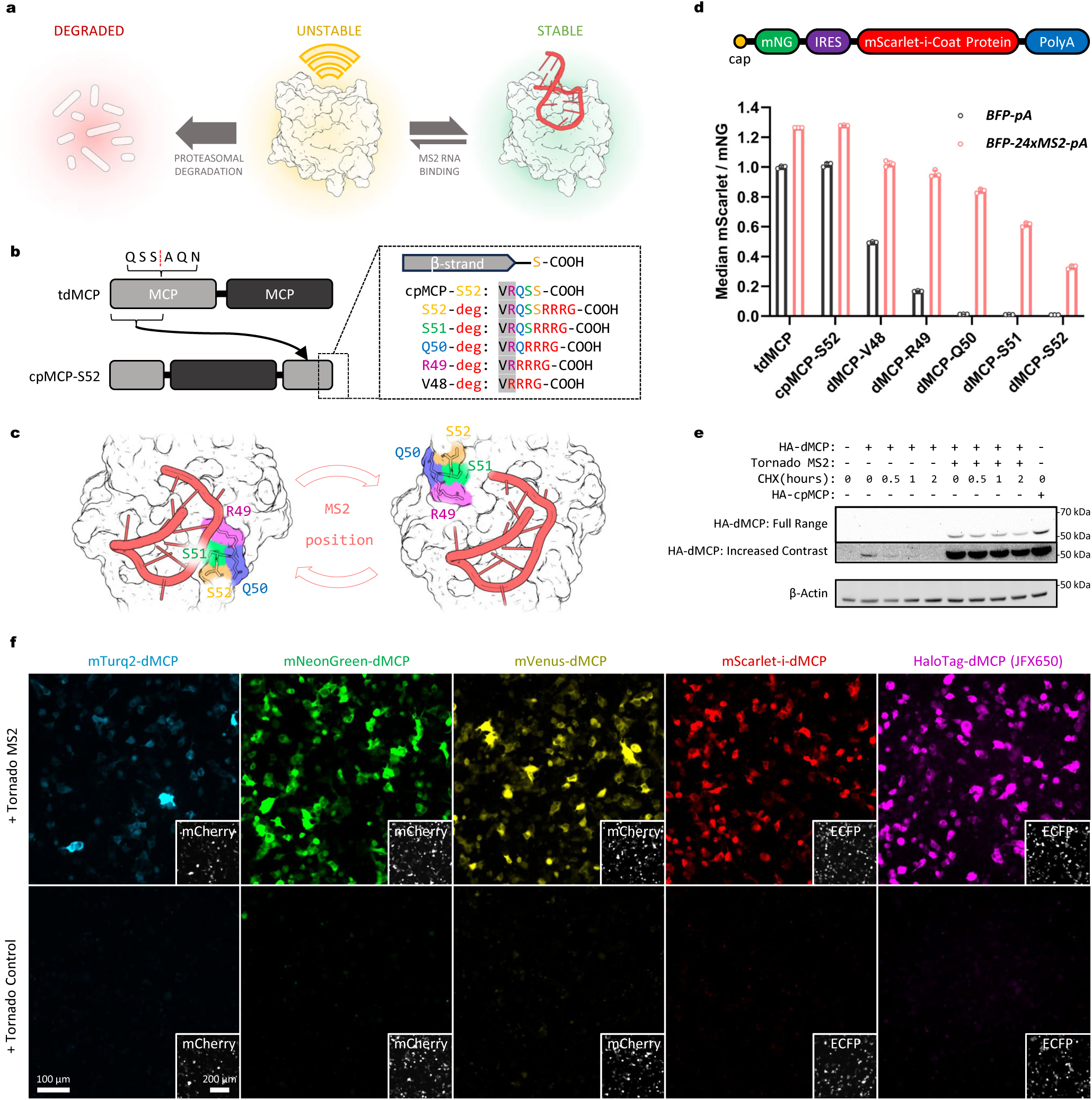
Design of an RNA-dependent conditionally stable MS2 coat protein. (**a**) Schematic depicting the RNA-dependent stability of an engineered MS2 coat protein. Unbound coat protein units are degraded within cells unless bound by MS2 hairpins. (**b**) Diagram showing the circular permutation strategy used on tdMCP. Attachment sites used to append the -RRRG C-degron to candidate sequences are also shown. (**c**) Structure of MCP showing the degron attachment sites indicated in (b) with juxtaposition to a bound MS2 hairpin RNA, as rendered using coordinates of a MS2-bound dimeric MCP complex (PDB: 1ZDH). (**d**) Design of the bicistronic gene construct used to quantify relative coat protein stabilities. Emissions from a mNeonGreen (mNG) reporter were used to normalize levels of IRES-driven mScarlet-coat proteins across cells and conditions. Below, mNG-normalized mScarlet levels are shown for the indicated tagged coat protein sequences, as measured via flow cytometry of transfected HEK-293FT cells co-expressing a *BFP-24xMS2* mRNA (red bars) or a *BFP* control mRNA (black bars). Individual points, bars, and error bars represent the individual median intensities per sample, the mean, and the S.D. of three independent transfections (n=3). (**e**) Cycloheximide chase of an mNG-fused HA-dMCP, as co-expressed in HEK293FT with either tornado-MS2 or MS2-lacking tornado-control. dMCP concentration and half-life under CHX treatment are visibly increased by the presence of MS2 RNA (**f**) Widefield fluorescence images of HEK293FT cells expressing the indicated dMCP fusions and RNAs. Insets depict the expression of a co-transfection marker. Emissions from fluorescent dMCP fusions were visible only in tornado-MS2 expressing cells.

Finally, using a similar approach, we also successfully generated a conditionally stable PCP. Equipped with two orthogonal and destabilized coat proteins, we exploited our new tools to simultaneously visualize distinct RNAs under low-background settings together in live cells. Overall, our approach combines the advantages of RNA-dependent signal generation, the versatility of genetic fusion proteins, and the well-characterized properties of RNA-binding coat proteins to enable sensitive and multicolor imaging of multiple RNA species.

## Design of a circularly permuted and destabilized MS2 coat protein

We first sought to create an MCP variant with a C-terminus that would become inaccessible upon MS2 hairpin binding. To generate such a variant, we used a tandem dimeric MCP (tdMCP)^21^ to create a circular permutant containing new termini within the RNA-adjacent EF loop (**Fig. 1b,c**). Since the resulting permutant terminated at the Ser52 position, we designated the construct “cpMCP-S52.” We confirmed that cpMCP-S52 retained its ability to bind MS2 RNA in mammalian cells by imaging the RNA-dependent relocalization of NLS-tagged cpMCP-S52 into the cytoplasm (**Supplementary Fig. 1a,b**).

Having verified its MS2-binding activity, we next appended a C-degron tetrapeptide (-RRRG)^20^ to cpMCP-S52 to produce a “destabilized MCP-S52,” or “dMCP-S52,” which we expected to degrade rapidly in cells without MS2 hairpins. Consistent with this expectation, dMCP-S52 levels were depleted compared to that of cpMCP-S52 when expressed in HEK293FT cells without tagged mRNAs (**Supplementary Fig. 1c**). In contrast, in cells containing MS2-tagged transcripts, dMCP-S52 levels were elevated. Together, these results indicate that the degron-mediated elimination of dMCP-S52 is curtailed to some degree in response to its specific binding to MS2-containing RNAs.

## Optimization of degron positioning

Previous work has shown that minor adjustments in C-degron positions, or changes in their relative flexibility and sequence contexts, can substantially affect their recognition by E3-ligases and, as a result, the degradability of their fused proteins.^22^ Thus, we asked whether the coat protein’s MS2-induced preservation could be optimized by adjusting the C-degron’s position on dMCP-S52. To evaluate this possibility, we varied the degron’s attachment position by deleting amino acids between the C-degron and the β-strand preceding it. These manipulations produced dMCP-S51, dMCP-Q50, dMCP-R49, and dMCP-V48, named based on the penultimate residues preceding their fused C-degrons and in which the degron tetrapeptide was positioned increasingly closer to the coat protein core (**Fig. 1b,c**).

To quantify the performance of these variants, we used bicistronic constructs to co-express a mNeonGreen (mNG) marker in combination with mScarlet-fused coat protein variants via an internal ribosome entry site (IRES). With this system, it is possible to normalize mScarlet-dMCP fluorescence across constructs and samples by comparing emissions from mNG. Flow cytometry was used to measure the relative expression of mScarlet-coat proteins in co-transfected HEK293FT cells, with comparisons between cells co-expressing an MS2-tagged mRNA (*BFP-24xMS2*) or an untagged transcript as a control (*BFP*).

Quantitative analyses of the variants revealed improved sequences, with the 1- and 2-amino acid deletion mutants exhibiting enhanced MS2-induced stability in *BFP-24xMS2* cells (**Fig. 1d**). Specifically, levels of MS2-preserved dMCP-Q50 were increased by 2.5-fold compared to that of the original dMCP-S52 design, with the deletion mutant maintaining minimal background levels in control cells. In *BFP-24xMS2-*containing cells, normalized dMCP-Q50 levels were elevated by an average of 63-fold compared to cells with the *BFP-pA* control. Further analysis of our flow cytometry data indicated a positive correlation between preserved coat protein levels and emissions from BFP, in which high BFP-expressing cells (above the 70^th^ percentile) exhibited a 100-fold enhancement in dMCP-Q50 levels in cells with *BFP-24xMS2* (**Supplemental Fig. 2b-d**). These data provide evidence for the MS2-specific preservation of dMCP-Q50, further suggesting a proportionality between stabilized dMCP-Q50 levels and the amounts of MS2-tagged mRNAs within cells.

Mutants with 3 and 4 deleted residues (dMCP-R49 and dMCP-V48, respectively) were also stabilized by MS2—however, basal background levels for these coat proteins were substantially elevated in *BFP* cells (**Fig. 1d**). As a result, these sequences had reduced ability to report on MS2-containing RNAs selectively.

Together, these results indicate that degron positioning is a critical and tunable parameter for producing conditionally stable RNA-binding coat proteins. Overall, we identified dMCP-Q50 as an optimal sequence, exhibiting marginal stability under basal conditions while being stabilized by up to two orders of magnitude in cells with MS2-containing RNA. Given these results, we thus proceeded with dMCP-Q50, hereafter designating it simply as ‘dMCP’ for ‘destabilized MCP’.

## MS2-binding extends the half-life of dMCP by inhibiting proteasomal degradation

With our design finalized, we next aimed to characterize the mechanism and dynamics of dMCP stabilization and degradation. First, we confirmed that dMCP’s low basal levels arise due to its active degradation by proteasomes. Indeed, fluorescence levels in cells expressing an mNG-fused dMCP were substantially elevated following treatment with the proteasome inhibitors lactacystin and MG-132 (**Supplementary Fig. 3**).

Next, we evaluated dMCP’s half-life in cells with and without MS2. Here, we co-expressed an HA-tagged dMCP (HA-dMCP) with long-lived circular RNAs,^23^ comparing a sequence containing a single inserted MS2 loop (MS2-tornado) against an MS2-lacking control (tornado-control). We used co-transfected HEK293-FT cells to conduct cycloheximide (CHX)-based chase analyses, and western blotting was applied to determine effective coat protein half-lives. Consistent with our prior results, these analyses showed that HA-dMCP accumulated in MS2-tornado cells to more than 50 times its levels in cells with tornado-control (**Fig. 1e**). As expected, the minimal HA-dMCP background in control cells was rapidly depleted upon CHX treatment, with the unliganded protein exhibiting an apparent half-life of less than 30 minutes (**Supplementary Fig. 4a,b**). By contrast, HA-dMCP was abundant in MS2-tornado cells, with levels that persisted during CHX treatment, suggesting a half-life of greater than 2 hours for the MS2-bound coat protein form. We also conducted pulse-chase measurements to evaluate dMCP’s stability without subjecting cells to global translation inhibition. Here, sequential dye labeling of dMCP fused with the self-labeling HaloTag^24^ (2xHalo-dMCP) indicated a half-life of approximately 10 hours in cells with MS2-tornado RNA (**Supplementary Fig. 4c,d**). Together, these results show that dMCP is rapidly eliminated by proteasomes and that its half-life is significantly extended upon binding MS2 hairpins within cells.

## Diverse dMCP-reporter fusions enable robust and selective RNA detection

To evaluate dMCP’s versatility as an RNA imaging probe, we tested dMCP constructs fused with different FPs and reporter domains. FPs from distinct lineages and of diverse colors were fused to dMCP’s N-terminus and co-expressed in cells with and without MS2-tornado. Imaging of co-transfected cells confirmed the conditional stability of the generated constructs, verifying dMCP’s compatibility with diverse FPs while also providing a multicolor toolset for imaging MS2-containing RNAs (**Fig. 1f, Supplementary Fig. 5a**).

We further tested dMCP using HaloTag and NanoLuciferase (NLuc) fusions. For cells expressing a HaloTag-dMCP, bright emissions from a JaneliaFluorX-650 (JFX650)-based ligand^25^ were observed only upon staining cells co-expressing MS2-tornado. (**Fig. 1f, Supplementary Fig. 5a**). The bioluminescent activity of NLuc-dMCP was also similarly MS2-responsive (**Supplementary Fig. 5b**). These results highlight dMCP’s versatility, confirming its compatibility with diverse reporters of distinct colors and modalities.

## dMCP enables high-contrast imaging of single mRNA molecules

Next, we evaluated the efficiency and accuracy of dMCP as a single mRNA probe. To obtain a construct of sufficient brightness for single-molecule imaging, we generated a 4xmNeonGreen-tagged dMCP (4xmNG-dMCP) and co-expressed the protein alongside an *H2B-mCherry-24xMS2* transcript in U2OS cells. Imaging of fixed cells by spinning disc confocal microscopy revealed distinct 4xmNG-dMCP intensities, which were visible in H2B-mCherry-positive cells with a signal-to-noise ratio (SNR) of 25 against cytoplasmic backgrounds (**Fig. 2a,b**, **Supplementary Fig. 6a-d**). To confirm that the visualized puncta accurately represented the locations of MS2-tagged transcripts, we used fluorescence *in situ* hybridization (FISH) to label fixed cells with antisense probes against *mCherry*. Following signal amplification by hybridization chain reaction (HCR)^26^, two color imaging revealed consistent co-registration between 4xmNG-dMCP puncta and emissions from AlexaFluor647 (AF647)-conjugated HCR probes. Quantification of these intensities showed that 92 ± 5% of the AF647 puncta also exhibited emissions from 4xmNG-dMCP (**Supplementary Fig. 6f**), with 4xmNG-dMCP labeling producing signals of relatively uniform dimensions and intensities, suggesting its consistent binding stoichiometry with MS2 arrays (**Fig. 2b,c Supplementary Fig. 6d,e**). We also used a 1xHaloTag-dMCP stained with a JFX650-based ligand in photobleaching spot analysis, calculating a ∼50% binding occupancy for the protein against a 16xMS2 array **(Supplementary Fig. 7)**. While lower than might be expected, this occupancy level consistent with values previously reported for tdMCP.^21^

**Figure 2.**
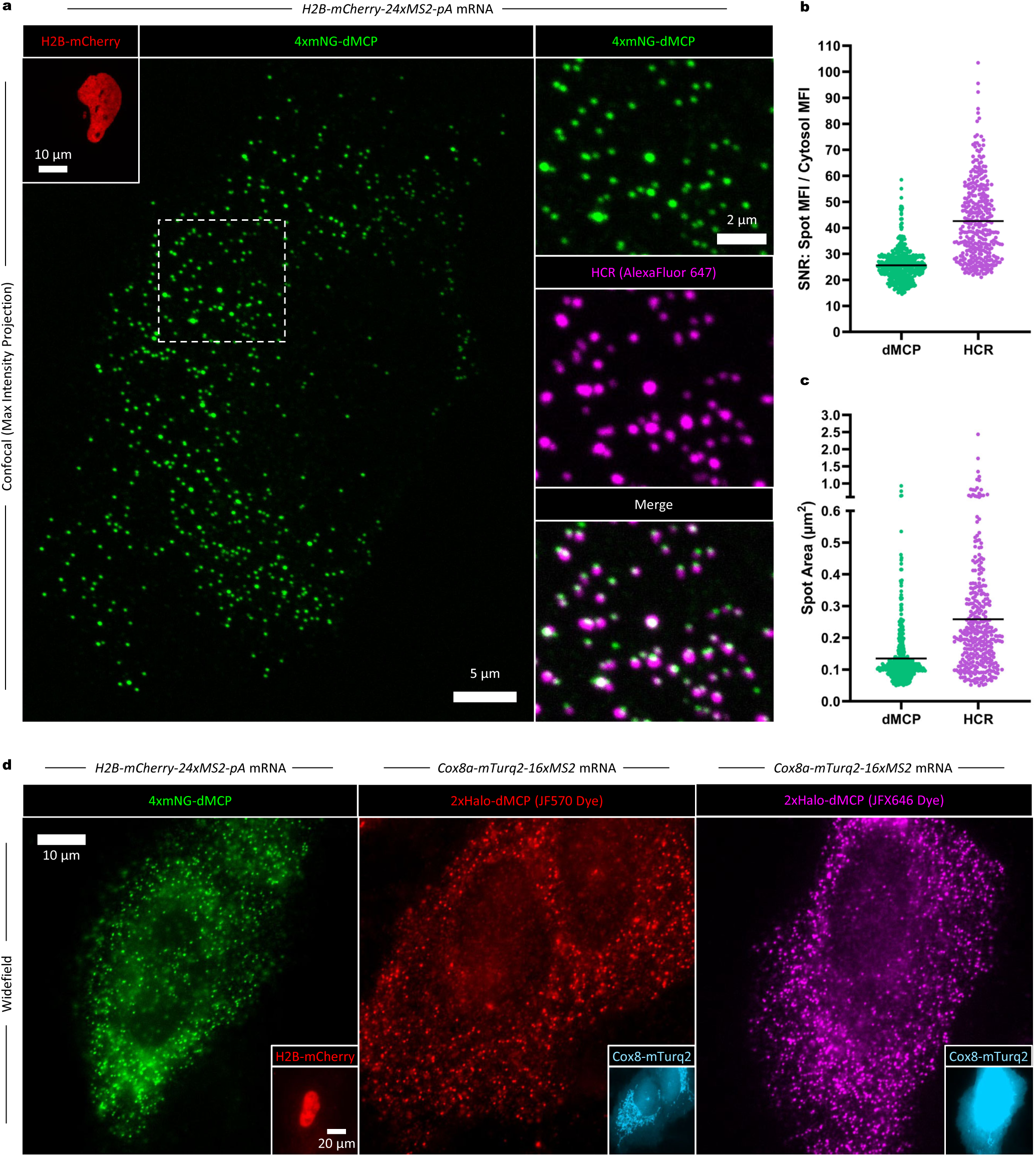
Single mRNA detection of MS2-tagged transcripts cells co-expressing 4xmNG-dMCP. (**a**) Images of a fixed U2OS cell expressing 4xmNG-dMCP and *H2B-mCherry-24xMS2-pA* transcripts. Co-localization of 4xmNG emissions with *mCherry* transcripts was confirmed via HCR-FISH using AF647-conjugated DNA probes. Maximum intensity projections from confocal microscopy are shown. (**b**) Signal-to-noise ratio (SNR), and (**c**) average size of individual 4xmNG-dMCP and HCR puncta within the imaged cell. SNR values in (b) are shown as mean intensities of individually measured punctae divided by the mean cytosolic intensity (excluding puncta). Black bars represent the mean for each population, with n=447 puncta for 4xmNG-dMCP and n=345 puncta for AF647. Larger HCR spot sizes resulted in reduced spot counts, such that certain individual AF647 puncta correspond to multiple 4xmNG-dMCP punctae. (**d**) Widefield images of fixed U2OS cells co-expressing the indicated dMCP fusions and MS2-tagged mRNAs. Insets represent detection of the protein fusions encoded by the corresponding MS2-tagged mRNAs.

To evaluate dMCP under more challenging imaging conditions, imaged cells using widefield microscopy, where minimized background fluorescence is essential for accurately detecting single molecules, given the expanded illumination depth across thicker z-planes. Under widefield imaging, 4xmNG-dMCP labeled transcript yielded distinct single-RNA signals within co-transfected cells (**Fig. 2d, left**). Single transcripts labeled with 2xHaloTag-dMCP were also distinctly visible upon staining with multiple fluorescent ligands, including those labeled with the photosensitizing (and thus rapidly photo-bleaching) JF570 chromophore,^27^ used previously in visualizing HaloTag fusions by correlative light and electron microscopy (**Fig. 2d, center**). Furthermore, staining 2xHaloTag-dMCP with the fluorogenic JFX646 enabled clear imaging of single transcripts via far-red emissions (**Fig 2d, right**)^25^.

To better image nuclear RNA, using lower mass fusion proteins, such as HaloTag-dMCP (‘1xHalo-dMCP’), is advantageous given its increased nuclear accessibility via passive translocation. Accordingly, imaging of JFX650-stained HaloTag-dMCP in HEK293FT cells revealed bright cytoplasmic and nuclear intensities, the latter of which likely reprsent nascent RNAs at sites of bursting transcription (**Supplementary Figure 8**). Together, these results validate dMCP as an accurate and sensitive probe for localizing MS2-tagged single mRNAs.

## Imaging the influence of nuclear retention elements on single RNA molecules using dMCP

Many RNAs are retained in the nucleus, where they can participate in processes such as splicing regulation, ribosome production, and transcriptional control, among other functions.^28,29^ Additionally, mRNAs with ‘detained introns’ can be blocked from nuclear export, with such blockage preventing the export and translation of incompletely spliced mRNAs as a mechanism for quality control.^30^ Under specific conditions, detained introns can also serve as regulatory elements, with certain detained sequences becoming spliced from pre-existing transcripts to facilitate their maturation and trigger their export in response to specific stimuli.^31^

Given their diverse roles, we thus used dMCP to selectively image nuclear RNAs, the visualization of which is challenging to do using conventional NLS-fused coat protein probes. In these analyses, we generated MS2-tagged transcripts by inserting the nuclear retention element (NRE) from the Maternally Expressed Gene 3 (MEG3) lncRNA (previously shown to facilitate the nuclear confinement of chimeric mRNAs)^32,33^ into the 5’ untranslated region (UTR) of *BFP-24xMS2* and *β-Globin-24xMS2* mRNAs. We then used a 1xmNG-dMCP to compare the localizations of NRE-containing and NRE-lacking transcripts in co-transfected HEK293FT cells. In these analyses, we observed strong nuclear retention of *MEG3NRE-β-Globin-24xMS2,* whereas the NRE-lacking *BFP-24xMS2* and *β-Globin-24xMS2* transcripts were predominantly cytoplasmic, both as expected (**Fig. 3a, Supplementary Fig.9**). Surprisingly, imaging of *MEG3NRE-BFP-24xMS2* revealed a predominately cytoplasmic localization, despite containing an inserted NRE (**Fig. 3a, Supplementary Fig.9**). This observation suggests that the nuclear retention of NRE-containing chimeric mRNAs may depend on specific sequence contexts, as suggested by previous work in which spliceosome-based components were identified as required factors in facilitating NRE-mediated mRNA retention.^32^ Thus, our data suggest that *MEG3NRE-BFP-24xMS2* is exported from the nucleus via a pathway that involves bypassing splicosomes, as potentially facilitated by its intronless nature.

**Figure 3.**
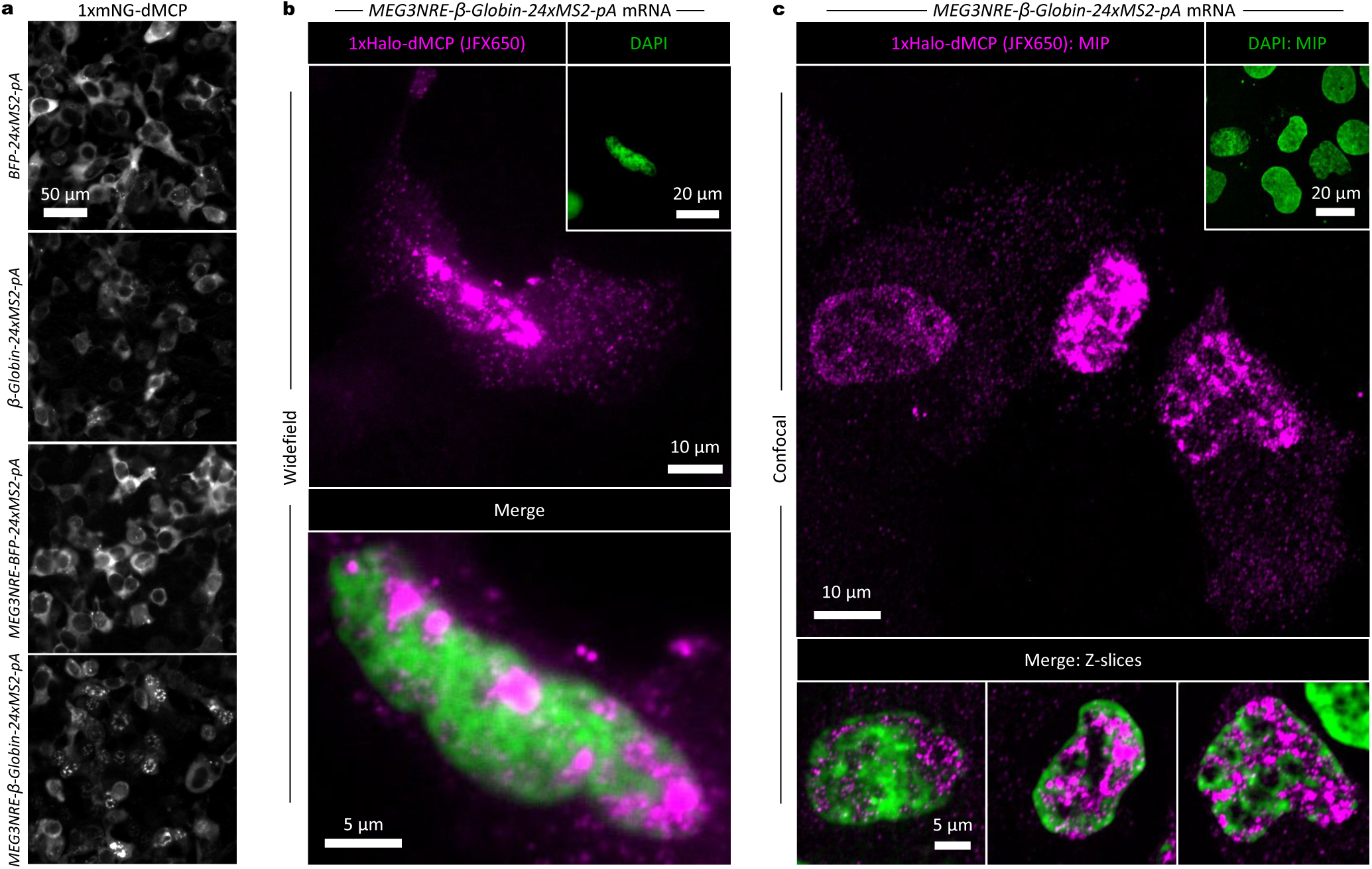
dMCP-based imaging nuclear retention element (NRE)-containing mRNA transcripts. (**a**) Widefield images of transfected HEK293FT cells co-expressing 1xmNG-dMCP in combination with the indicated MS2-tagged mRNAs. NRE-containing and NRE-lacking transcripts encoding BFP or β-Globin are compared. (**b**) Widefield fluorescence images of a U2OS cell co-expressing 1xHaloTag-HA-dMCP stained with JFX650 (magenta) and *MEG3NRE-β-Globin-24xMS2-pA.* Nuclear DNA was visualized via DAPI staining (green). (**c**) Confocal images of cells prepared as in (b). NRE-containing mRNAs are seen accumulated within DNA-depleted interchromatin regions (areas exhibiting reduced DAPI emissions), and a subset of tagged mRNA are visible in the cytoplasm. Single-channel confocal images are maximum intensity projections; and confocal overlay images are single z-slices.

We next used dMCP to visualize the distribution of single NRE-containing transcripts with subnuclear detail. Previous work has shown that MEG3-NRE insertion causes mRNAs to associate with nuclear speckles,^33^ subnuclear components that are rich in RNA and protein, while containing limited amounts of DNA. To image such associations, we used HaloTag-dMCP in combination with a JFX650-based ligand to visualize single *MEG3NRE-β-Globin-24xMS2* mRNAs under widefield and spinning disc confocal imaging (**Fig. 3b-c**). To provide nuclear context, we visualized the labeled mRNAs in cells stained with 4’,6-diamidino-2-phenylindole (DAPI), reasoning that by juxtaposing the tagged transcripts with DAPI emissions, we could identify nuclear speckle-associated transcripts based on their existence within DAPI-negative nuclear areas. Indeed, high-magnification imaging revealed enrichment of labeled transcripts within chromatin-lacking subnuclear locales, agreeing with their expected association within DNA-depleted interchromatin granule clusters.

## dMCP enables versatile and sensitive live cell imaging of single mRNA molecules

Having confirmed dMCP as a sensitive RNA probe in fixed cells, we next evaluated its utility in imaging the live cell dynamics of single mRNAs. Using U2OS cells co-expressing an *H2B-mCherry-16xMS2* transcript in combination with 4xmNG-dMCP, we used widefield microscopy to record the trajectories of tagged mRNAs in real-time. Recorded movies revealed dMCP-labeled transcripts as bright intensities undergoing diffusive Brownian motions within the U2OS cytoplasm (**Supplementary Movie 1**). Analysis of these trajectories showed that a majority of transcripts remained visible over the recorded durations, likely due to the broad imaging depth of widefield microscopy, which, unlike confocal imaging, allows fluorescent particles to remain visible as they traverse distances along the z-axis dimension.

To investigate the relationship between the observed diffusive motions and translation activity, we imaged cells before and after treatment with the translation inhibitor harringtonine (HT). Apparent diffusion coefficients (D_app_) of single *H2B-mCherry-16xMS2* transcripts were significantly increased upon HT treatment, consistent with their expected mass reductions due to HT-mediated polysome dissociation (**Fig. 4a,b, Supplementary Figure 10a,b, Supplementary Movie 1**). This result agrees with previous TIRF imaging studies, where translating transcripts moved more slowly on average than transcripts under translational repression by puromycin.^34^ Mean Squared Displacement (MSD) analysis produced a calculated diffusion coefficient of 0.13 µm^2^/s for dMCP-tagged *H2B-mCherry-16xMS2* mRNA under normal conditions (**Supplementary Figure 10c,d**), in line with previously reported values for MS2-tagged mRNAs labeled by conventional MCP.^14,35^ Seeking to confirm its live imaging utility in multiple cellular contexts, we also visualized single mRNAs in primary neonatal human dermal fibroblasts (HDF-Neo), applying lentiviral transduction to co-express a 2xmNG-dMCP in combination with an *ECFP-24xMS2-pA* transcript in these cells (**Supplementary Movie 2**). Together, these data highlight dMCP’s utility in tracking single mRNAs across whole cell volumes, including in primary cells.

**Figure 4.**
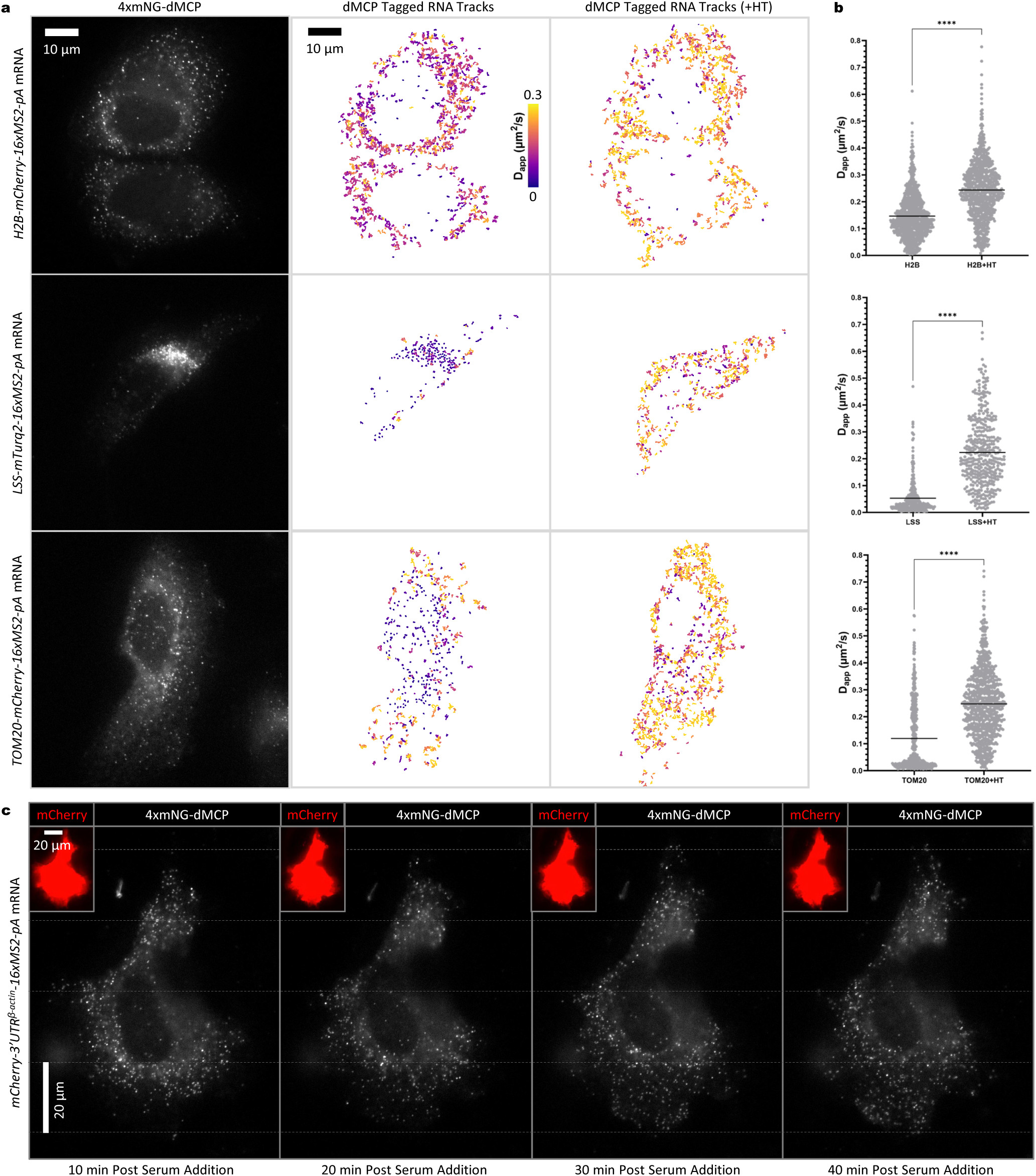
Live-cell tracking of MS2-tagged single mRNAs in 4xmNG-dMCP-expressing U2OS cells. (**a**) Single-frame captures from video recordings of 4xmNG-dMCP emissions in cells expressing the indicated 16xMS2-tagged mRNAs (left). RNA trajectories from recordings are shown as color-graded single mRNA particle traces before (middle) and after (right) harringtonine (HT)-induced translation inhibition. Only traces lasting at least 3 frames (0.26 seconds at 0.085 sec/frame) are displayed and analyzed. Traces are color-graded based on apparent diffusion coefficients, calculated using mean squared displacement (MSD) values with a time delay of 1 frame. (**b**) Apparent diffusion coefficients for the tracked single RNA before and after HT treatment. Statistical significances were determined via unpaired two-tailed Student’s t-test: ****P<0.0001. Trace counts: H2B n=899, H2B +HT n=814, LSS n=295, LSS +HT n=401, TOM20 n=519, TOM20 +HT n=829. (**c**) Images of a U2OS cell containing 4xmNG-dMCP labeled mCherry-encoding transcripts bearing an inserted 3’UTR^β-actin^. Redistribution of transcripts towards cell edges can be observed following serum stimulation of serum-starved cells. Insets display saturated mCherry emissions to highlight cell boundaries.

Next, we used dMCP to investigate differential mRNA dynamics for transcripts that encode organelle-targeted or secreted model proteins. We first evaluated an mRNA encoding a secreted protein containing the lysozyme secretion signal (*LSS-mTurq2-16xMS2*). In these analyses, labeled mRNAs were visualized as static intensities, consistent with their expected docking on the surface of the rough ER (**Supplementary Movie 3**). Treatment with HT converted the static puncta into rapidly diffusing intensities (**Fig. 4a,b, Supplementary Fig. 11, Supplementary Movie 4**), with the transcripts returning to their static states following HT washout (**Supplementary Fig. 12**). Collectively, these results provide a clear and direct visualization of the well-characterized process co-translational secretion, during which ribosome-bound mRNAs dock to translocon complexes as they undergo translational elongation.^3^

We also applied dMCP to visualize mRNAs encoding a protein targeted to mitochondria via the N-terminal mitochondrial targeting sequence (MTS) from the Tom20 subunit of the mitochondrial outer-membrane translocase (TOM) complex (*MTS^Tom20^-mCherry-16xMS2*). Recording of 4xmNG-dMCP-labeled mRNAs revealed distinct transcript populations, including highly mobile and relatively immobile particle populations (**Supplementary Movie 5**). Spatial mapping of particle velocities showed distinct subcellular localizations for these populations, with the slow-moving transcripts predominantly distributed perinuclearly and often co-localizing with TOM20-mCherry labeled mitochondria (**Supplementary Fig. 13**). Treatment with HT resulted in the conversion of *MTS^Tom20^-mCherry-16xMS2* transcripts into distributed and freely diffusing intensities, indicating that their prior static states were dependent on active translation elongation (**Fig. 4a,b, Supplementary Fig. 14, Supplementary Movie 6**). These results suggest that a subset of mRNAs encoding proteins with TOM20’s 33-amino acid MTS may transiently associate with mitochondria via a translation-dependent mechanism, possibly through interactions between nascent targeting peptides and MIM1 complexes on mitochondrial outer surfaces.^36^

To test whether dMCP could visualize transcripts expressed at near-native levels, we used 4xmNG-dMCP to visualize CRISPR-tagged *TOMM20* mRNAs using knock-in HEK293FT cells bearing an inserted *mCherry-10xMS2* cassette at the endogenous TOM20-encoding locus (**Supplementary Fig 15**).^37^ Tagged transcripts in knock-in cells were clearly visible through transient 4xmNG-dMCP expression, with a fraction of labeled intensities appearing to co-localize with TOM20-mCherry mitochondria, agreeing with our observations using co-transfected U2OS cells (**Supplementary Movie 7**).

Finally, in addition to co-translational regulatory mechanisms, many mRNAs are targeted to subcellular locations through direct transport via RNA-based sorting signals within 3’-UTRs. A well-characterized example of such transport is that of β-actin transcripts, which contain a 3’ element (3’-UTR^β-actin^) that is recognized by zipcode binding protein-1 (ZBP-1, also called IGF2BP1)^38^ to facilitate directed mRNA transport along cytoskeletal filaments^39^. To visualize such trafficking, we used dMCP to track a *mCherry* transcript containing and inserted 3’UTR^β-actin^ signal upstream of an MS2 array (*mCherry-3’UTRβ-actin-16xMS2).* In cells co-expressing 4xmNG-dMCP, we observed the previously characterized effect of serum-induced redistribution of the 3’UTR^β-actin^-containing transcript to distal cellular regions^40^ (**Fig. 4c, Supplementary Fig. 16**). Together, these data highlight the utility of dMCP in studying the dynamics and regulated targeting of mRNAs across subcellular regions, including those that undergo targeting via translation-dependent and -independent processes.

## Generation of a destabilized PP7 coat protein (dPCP) for two-color imaging

Given our success in developing dMCP, we next asked whether we could design a destabilized PP7 coat protein, which orthogonally binds a cognate PP7 RNA hairpin.^41^ Here, we reasoned that by creating a destabilized PCP (dPCP), we could simultaneously visualize distinctly tagged transcripts in the same cell.

Structural analyses have shown that PCP and MCP share similar folds and RNA-binding characteristics despite limited sequence conservation,^8^ with PCP also orienting its EF loop proximally to PP7 in its hairpin-bound form. Given this orientation, we generated a circularly permuted tdPCP^21^ with new termini at the RNA-adjacent EF loop (**Fig. 5a**). Initial tests of the resulting permutant, cpPCP-G48, confirmed its binding and nuclear co-export with a PP7-containing tornado RNA (tornado-PP7). However, imaging of cpPCP-G48 also revealed visible protein aggregates formed in both tornado-PP7 and control cells (**Supplementary Fig. 17**). Hypothesizing that these aggregates may be due to homomeric oligomer formation, we thus introduced mutations to disrupt cis-interactions between cpPCP-G48 units. Such mutations produced a ‘solubilized cpPCP-G48,’ or ‘sol-cpPCP-G48,’ which lacked visible aggregation in transfected HEK293FT cells while retaining PP7-binding activity (**Supplementary Fig. 17**).

**Figure 5.**
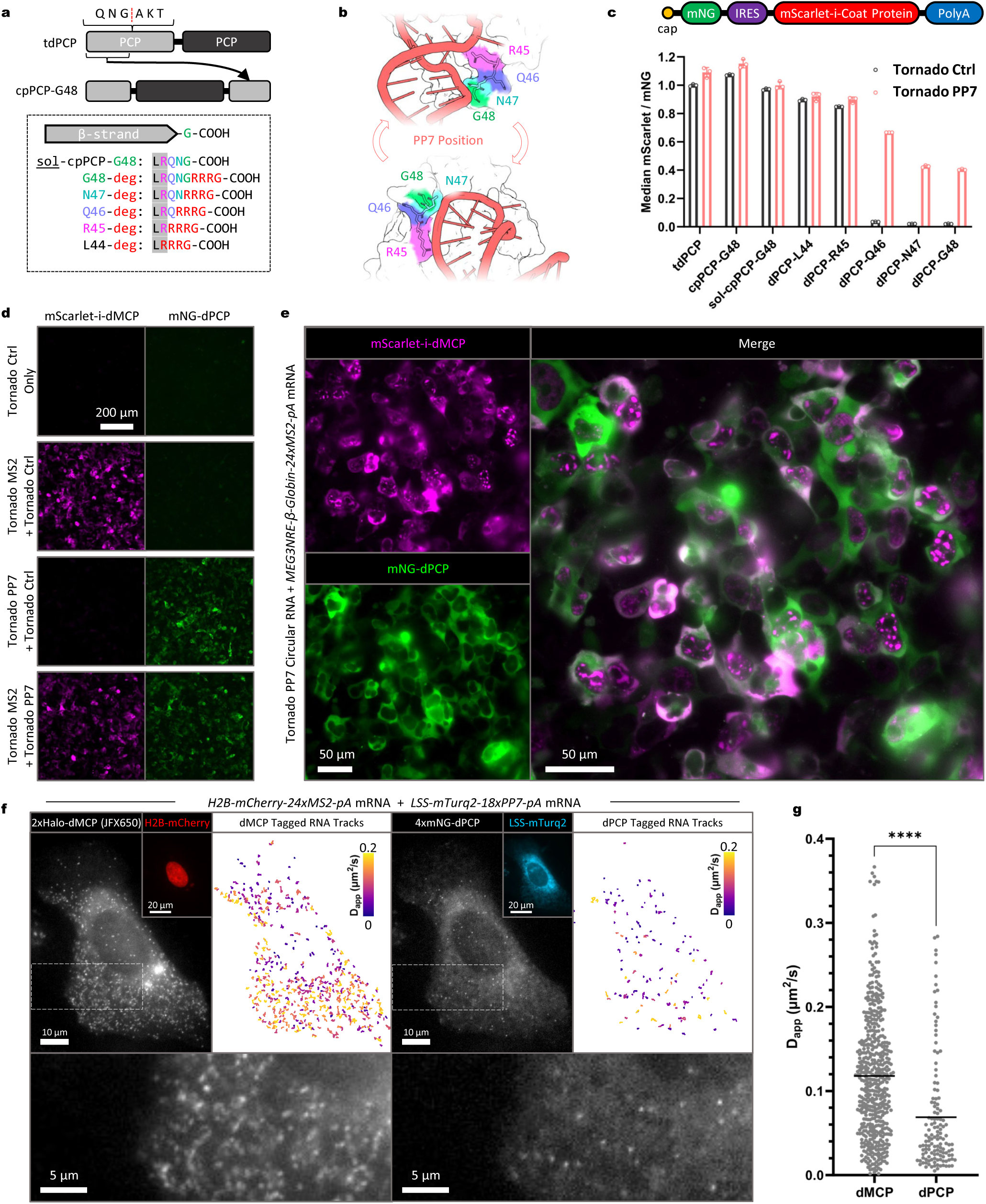
Design of destabilized PP7 coat protein and two-color RNA imaging using dMCP and dPCP. (**a**) Schematic showing the circular permutation strategy used to design dPCP; tdPCP permutation sites and degron attachment locations are displayed. A solubilized tdPCP circular permutant (“sol-cpPCP-G48”) was used to generate the candidate degron-tagged proteins. (**b**) Structures depicting the degron attachment positions as indicated in (a) and rendered using coordinates from native PP7-bound PCP (PDB: 2QUX). (**d**) Schematic of the bicistronic gene construct used to quantify relative degron-tagged permutant stabilities (top). Analyses were done in a similar manner as with dMCP, with use of mNG emissions to normalize levels of IRES-driven mScarlet-coat proteins fusions (bottom) in cells co-expressing tornado-PP7 (red bars) or tornado-control (black bars). Individual points, bars, and error bars represent the individual median intensities per sample, the mean, and the S.D. of three independent transfections (n=3). (**d**) Widefield images of HEK293FT cells co-expressing mScarlet-dMCP and mNG-dPCP with the indicated MS2-tagged, PP7-tagged, or untagged circular ‘tornado’ RNAs. (**e**) Distinct subcellular distributions of MS2- and PP7-tagged RNAs in transfected HEK293FT cells. Nuclearly-retained MS2 mRNA (*MEG3NRE-β-Globin-24xMS2-pA*) is labeled by mScarlet-dMCP (magenta), in juxtaposition to nuclearly-exported tornado-PP7 RNA labeled by mNG-dPCP (green). (**f**) Live cell multi-channel tracking of orthogonally tagged single mRNAs in a U2OS cell. Single transcripts corresponding to *H2B-mCherry-24xMS2-pA* (left) and *LSS-mTurq2-18xPP7-pA* (right) were detected via 2xHalo-dMCP labeled by JFX650 dye and 4xmNG-dPCP, respectively. Detection of the independently tagged transcripts are shown in grayscale as single frame captures; insets represent detection of the corresponding transcript-encoded proteins. Bottom panels represent magnified views of the areas within the dashed lines. Top right panels represent single RNA trajectories as color-graded traces. Traces lasting at least 3 frames (0.68 seconds at 0.226 seconds/frame) are shown. Color scales depict the apparent diffusion coefficients, as calculated using mean squared displacement (MSD) with time delays of 1 frame. (**g**) Calculated apparent diffusion coefficients for the indicated RNA transcripts. Statistical significances were determined by unpaired two-tailed Student’s t-test: ****P<0.0001. Trace counts: dMCP/MS2 n=507, dPCP/PP7 n=132.

Candidate destabilized sequences were generated by fusing the -RRRG C-degron to sol-cpPCP-G48, with a variation of the attachment positioning as before, producing dPCP-G48, dPCP-N47, dPCP-Q46, dPCP-R45, and dPCP-L44 (**Fig. 5a,b**). We then used a similar bicistronic vector to assay the stability of these sequences in cells with and without tornado-PP7 RNA, leading to the identification of dPCP-Q47 as an optimal variant, which exhibited a 19-fold stability enhancement in cells co-expressing tornado-PP7 RNA (**Fig. 5c, Supplementary Fig. 18**). Thus, we proceeded with dPCP-Q47, designating it hereafter simply as “dPCP.”

With dMCP and dPCP in hand, we sought to confirm their orthogonal ligand-induced stabilization and to combine these proteins to image two distinct RNA species in living cells simultaneously. As expected, dMCP and dPCP were selectively stabilized by their cognate hairpin ligands in co-transfected HEK293FT cells without observable cross-talk, confirming their maintained orthogonality (**Fig. 5d**). To exploit their orthogonality, we used the coat proteins to image the nuclear retention of a *MEG3NRE-β-Globin-24xMS2* transcript in cells alongside the cytosolically-targeted tornado-PP7, observing distinctly localized emissions from the associated coat proteins as expected (**Fig. 5e**).

Finally, we used 4xmNG-dPCP to resolve an *H2B-mCherry-18xPP7* mRNA with single molecule sensitivity, observing freely diffusing intensities consistent with those visualized using dMCP (**Supplementary Movie 8**). To further exploit their orthogonality, we combined a 4xmNG-dPCP and a 2xHaloTag-dMCP to simultaneously visualize *H2B-mCherry-24xMS2* and *LSS-mTurq2-18xPP7* mRNAs, observing both highly mobile *H2B-mCherry-24xMS2* and ER-bound *LSS-mTurq2-18xPP7* transcripts by widefield imaging, both as expected (**Fig. 5f,g, Supplementary Movie 9**). These data demonstrate the utility of dMCP and dPCP as robust and sensitive tools to visualize tagged RNAs independently and in combination via multicolor imaging.

## DISCUSSION

In summary, we developed MS2 and PP7 coat proteins with RNA-dependent stability, reducing background signals arising from RNA-unbound coat protein units. We achieved this through a protein engineering scheme in which we first applied circular permutation to position protein termini to RNA ligand-adjacent sites, which we then exploited as attachment sites for an RNA-maskable C-terminal degron. With this approach, we generated destabilized coat proteins that rapidly degraded in cells unless bound to their cognate RNA motifs. We demonstrated the destabilized coat proteins’ ability to detect and visualize distinctly localized single RNA molecules in live cells, including nuclear transcripts, with the lowered background permitting sensitive recording of single mRNA dynamics via widefield fluorescence imaging. Further, by developing two orthogonal protein-RNA pairs, we could detect and track distinct mRNA species together in cells simultaneously.

To create dMCP and dPCP, we implemented a C-degron insertion strategy that builds on previously implemented techniques.^18^ As the native ends of these coat proteins are located at non-maskable positions, we used circular permutation to relocate their C-termini to RNA-adjacent locations. By fusing an -RRRG degron to the permutants and screening positional variants, we created constructs with optimal RNA-dependent stabilities. Characterization of dMCP showed RNA-induced stabilization led to a 63-fold increase in intracellular concentration, with the coat protein becoming stabilized via a mechanism that extends its half-life within cells. We further demonstrated the robustness of this strategy by developing dPCP in an analogous way. However, preliminary protein engineering was required to increase the solubility of PCP-based sequences. Additionally, we note that dPCP’s performance is modest compared to that of dMCP (with dPCP exhibiting ∼19-fold enhancement in response to PP7 RNA). Despite its reduced stabilization, dPCP was suitable for visualizing single mRNAs, both independently and in combination with dMCP via two-color imaging.

Using dMCP and dPCP, we were able to track the localization and live cell dynamics of diverse RNAs via widefield imaging. Such RNAs included circular RNAs, nuclearly-retained transcripts, and single mRNAs encoding secreted, mitochondrial, or locally translated proteins. Experiments in CRISPR-modified lines and transduced primary cells show that dMCP will be useful in imaging RNA under varying cellular contexts, which we anticipate will facilitate its adoption in future biological studies. By using dMCP and dPCP together, we showed that two differentially regulated transcripts could be simultaneously tracked within the same cells. Overall, we expect dMCP and dPCP to be readily combined with existing systems utilizing MS2 and/or PP7 RNA motifs.

Limitations of dMCP and dPCP should be noted. Future users should be mindful of potential cellular effects arising from the expression of these proteins, which may occur due to their competition against endogenous degradation substrates. To minimize such burdens, we recommend expressing dMCP and dPCP fusions at low levels by using weak promoters or IRES-mediated expression or introducing only limited gene copies to cells. Such efforts may serve to dually minimize cellular effects while also ensuring the maintenance of low background levels. Additionally, since the background-suppressing degradation mechanism is a kinetic process, users should consider the stability and half-life of target RNA when applying these probes, as short-lived RNA transcripts may degrade too quickly for effective target binding, stabilization, and chromophore maturation.

All results presented herein were obtained in human-derived cell lines or primary cells, but the -RRRG degron we used has demonstrated effectiveness across a range of animal cell types, including various mammalian cells^42^ and *in vivo* within flies,^43^ zebrafish,^44^ and mice.^45^ The consistent performance of this specific degron across animal cells and models is likely due to the evolutionary conservation of E3 ligases that recognize this motif.^46^ However, dMCP and dPCP are not predicted to function in prokaryotes or fungi, which lack the specific degradation machinery necessary to recognize the -RRRG degron.^46^ Given that both termini of the circularly permuted variants are positioned adjacent to their RNA-binding sites, it may be feasible to screen additional C- or N-terminal degrons to develop dMCP and dPCP variants functional in other kingdoms and domains of life.

While we focused on applications of dMCP and dPCP for fixed and live-cell RNA imaging, we anticipate that these conditionally stable proteins could enhance performance for many RNA-based systems. In additional imaging applications, dMCP and dPCP could serve as high-contrast agents for genomic loci imaging using dCas9-sgRNA scaffolds.^47–49^ RNA adaptor systems may also enable the detection of endogenous RNA expression through dMCP and dPCP.^50^ Additionally, dMCP and dPCP may help reduce off-target background in proximity labeling and RNA-protein interaction detection applications.^51,52^ Finally, pairing dMCP or dPCP with effector domains, such as RNA^53–57^ or DNA^58,59^ modifying enzymes, could improve the on-target performance while minimizing nonspecific targeting of bystander genes. Often, these applications already use MS2 or PP7 as their RNA handle, and thus dMCP and dPCP are anticipated to be directly adaptable to these systems in most animal cell types.

## DATA AVAILABILITY

The datasets generated during and/or analyzed during the current study are available from the corresponding author upon reasonable request. Plasmid DNA and detailed sequence information for described constructs will be made available through AddGene upon acceptance of the publication.

## CONTRIBUTIONS

All authors contributed to the design and execution of experiments, analysis of the results, and preparation and editing of the manuscript.

### ACKNOWLEDGEMENTS

This work was funded through NIH research grant R35GM128859 (to J.T.N.). C.J.K. was supported through a Multicellular Design Program Fellowship (Boston University Kilachand Fund). A.M.M. was a recipient of a National Science Foundation Graduate Research Fellowship.

