## Supplementary Information for "RNA-Stabilized Coat Proteins for Sensitive and Simultaneous Imaging of Distinct Single mRNAs in Live Cells"

**CONTENTS:**

I. Supplementary Figures 1-18
II. Materials and Methods
III. Captions for Supplementary Movies 1-9
IV. Amino Acid Sequences for the Reported Protein Domains

### Supplementary Figure 1

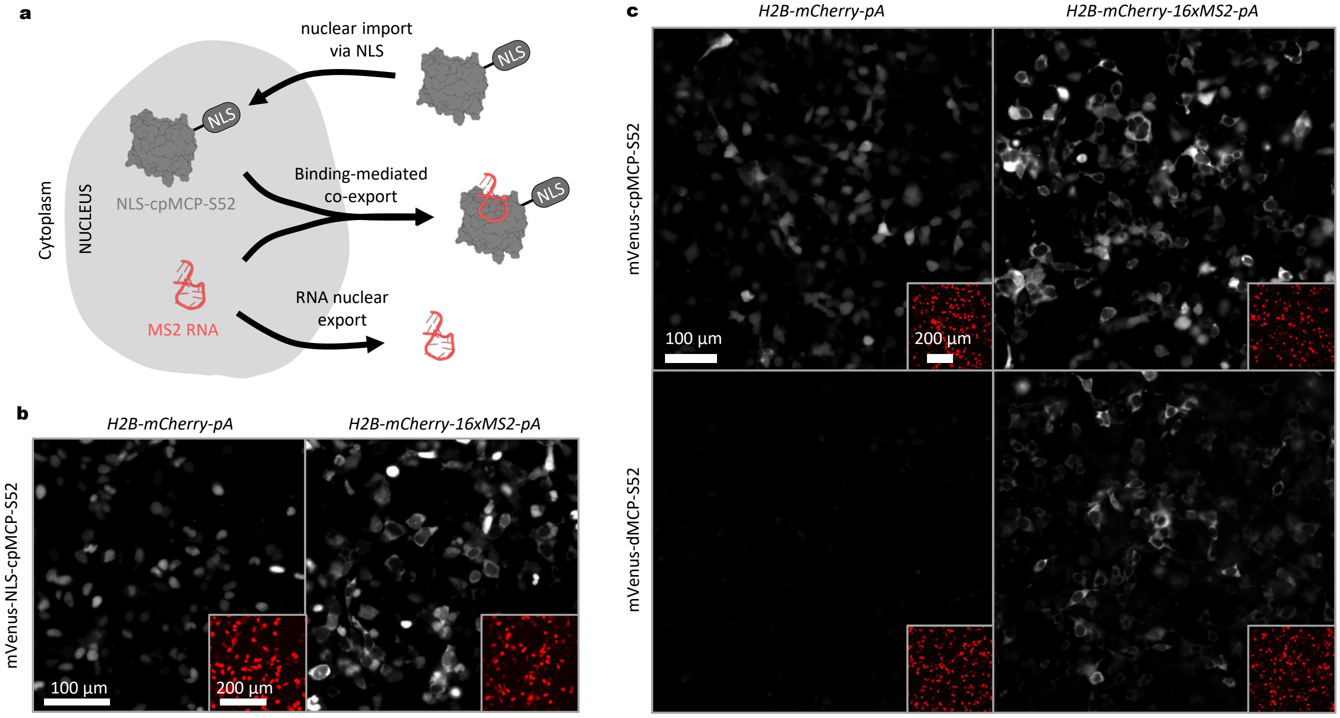

**Supplementary Figure 1. MS2-binding analysis using NLS-cpMCP-S52. (a)** Schematic depicting the RNA binding-mediated nuclear export of NLS-cpMCP-S52, which is imported into and retained within the nucleus in its unbound form. Upon overexpression and nuclear export of MS2-tagged mRNA, protein-RNA binding leads to translocation of NLS-cpMCP-S52 to the cytoplasm. **(b)** Widefield fluorescence microscopy images of HEK293FT cells co-transfected to express mVenus-NLS-cpMCP-S52 and either *H2B-mCherry-pA* or *H2B-mCherry-16xMS2-pA* transcripts show that expression of MS2-tagged mRNA leads to nuclear exclusion of NLS-cpMCP-S52, indicating that it is capable of binding to MS2 loops. Insets in the bottom right display H2B-mCherry expression **(c)** HEK293FT cells were transfected to express mVenus-cpMCP-S52 or the degron-appended mVenus-dMCP-S52 (both lacking an NLS). These cells were cotransfected to express *H2B-mCherry-pA* or *H2B-mCherry-16xMS2-pA* transcripts. The images show that the fluorescence of mVenus-dMCP-S52 is heavily reduced in the absence of MS2 RNA, and that this fluorescence is partially recovered in the presence of MS2 RNA.

### Supplementary Figure 2

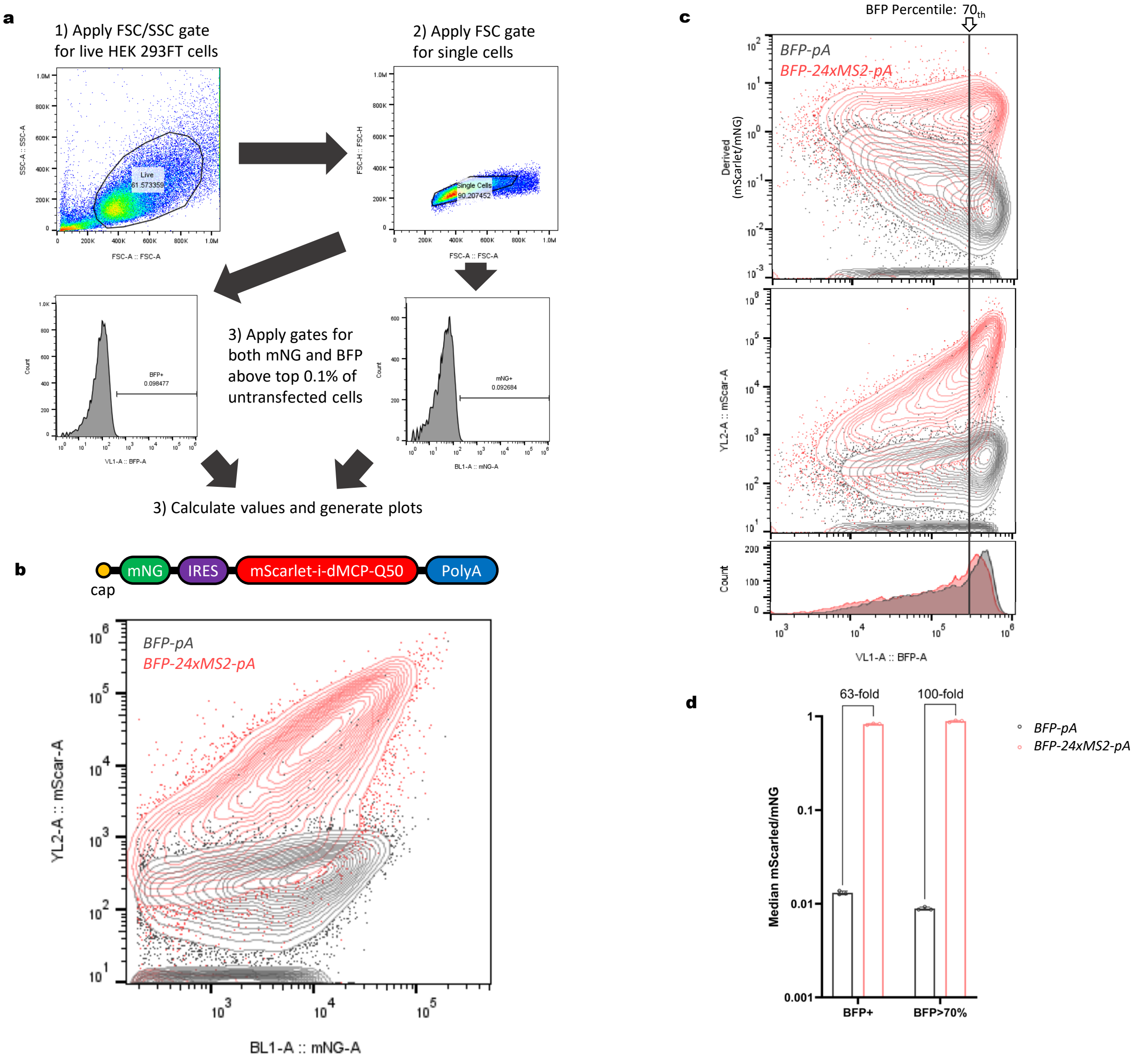

**Supplementary Figure 2. Flow cytometry gating scheme and results of different degron integrated cpMCP variants. (a)** Flow cytometry gating method. Cells were first gated for live singlets based on forward and side scatter. Cells were then gated for BFP and mNeonGreen reporter expression above the top 0.1% of untransfected cells. **(b)** Schematic of the reporter construct, with cap-dependent translation of a stable mNeonGreen expression reporter and IRES-dependent translation of the optimal dMCP-Q50 fused to mScarlet-i. Beneath, representative flow cytometry data from HEK293FT cells expressing the above construct and *BFP-pA* (gray) or *BFP-24xMS2-pA* (red). mScarlet intensity is plotted versus mNeonGreen intensity to display the relation between dMCP expression level and its response to the presence of MS2-RNA. **(c)** Representative flow cytometry data of the dMCP-Q50 response to increasing concentrations of MS2 RNA (as indicated by BFP expression). In the top plot, mNeonGreen-normalized mScarlet-dMCP intensity is plotted. In the middle plot, absolute mScarlet-dMCP intensity is plotted. The bottom histogram indicates the distribution of BFP expression within mNeonGreen positive cells, with 70$^{th}$ percentile marker overlaid across all three graphs. **(d)** Median mScarlet-dMCP intensities for the entire BFP positive population and the top 30% of BFP expressing cells. Quantification performed by flow cytometry in HEK293FT cells co-transfected with BFP expression vectors with (red circles) or without (black circles) MS2-arrays. Each point represents the median normalized mScarlet expression for independent transfections. Bars and error bars represent the mean and S.D of three independent transfections (n=3).

### Supplementary Figure 3

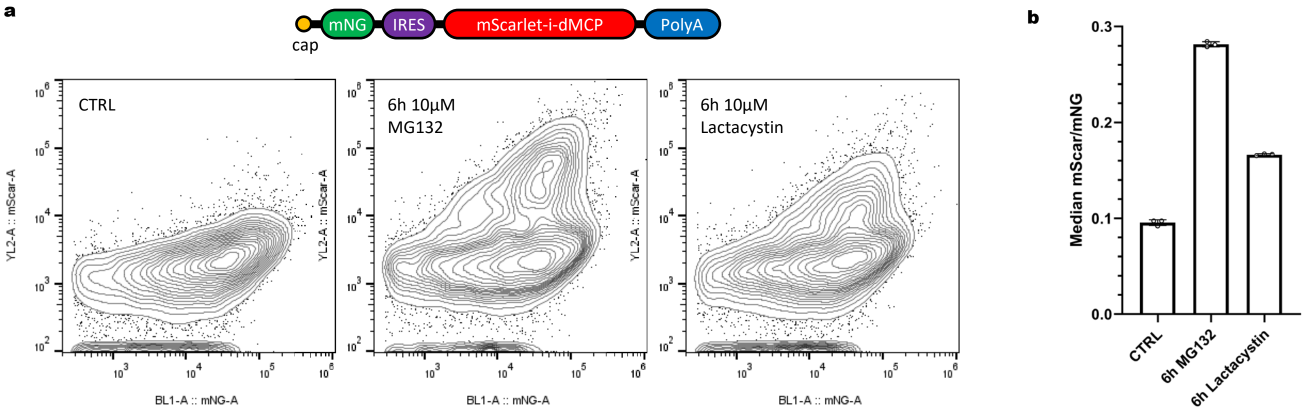

**Supplementary Figure 3. Low basal dMCP expression is partially rescued by proteasome inhibition. (a)** Representative flow cytometry traces of HEK293FT cells transfected with the bicistronic mNeonGreen-IRES-mScarlet-dMCP-Q50 construct under standard conditions (left) and after 6 hours of proteasome inhibition with 10 μM MG132 (middle) or 10 μM lactacystin (right). A partial increase in the mScarlet intensity of a subset of the population can be observed under proteasome inhibition. **(b)** Quantification was performed by flow cytometry for each population. Each point represents the median normalized mScarlet expression for an independent transfection. Bars and error bars represent the mean and S.D of three independent transfections (n=3).

#### Supplementary Figure 4

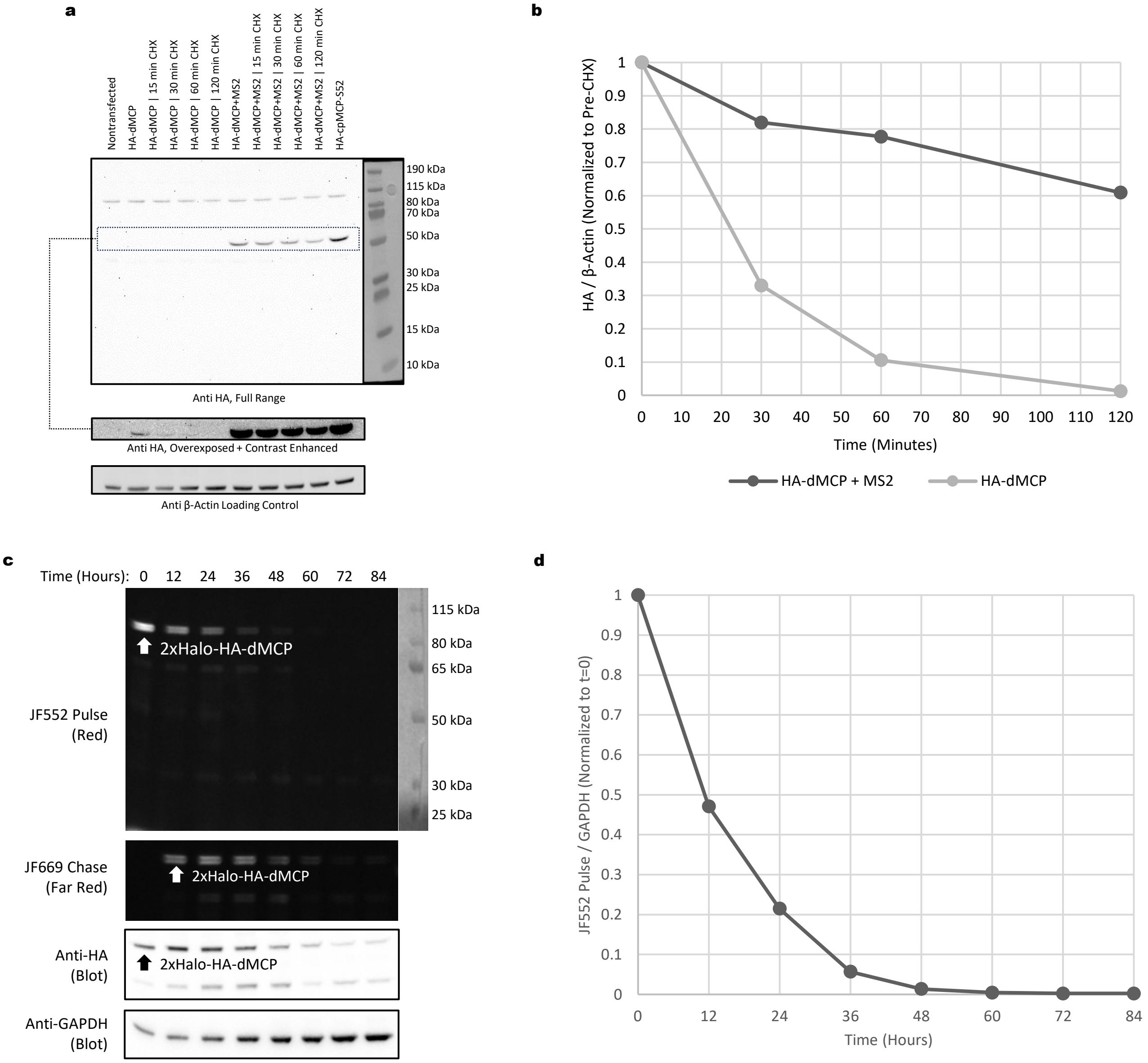

**Supplementary Figure 4 - Full membranes and gels of dMCP cycloheximide chase experiment and HaloTag pulse chase experiment. (a)** Full nitrocellulose membranes corresponding to the cycloheximide (CHX) chase data shown in Fig 1e. Overexposed HA-stain to resolve unstable dMCP are shown below along with a β-actin loading control. ImageJ was used to quantify band intensities, and a 50-fold difference was measured for β-actin-normalized HA-dMCP band intensities between tornado MS2 and tornado control conditions prior to cycloheximide treatment **(b)** Plotted normalized HA-dMCP band intensities in CHX-treated cells over time shows that dMCP half-life is less than 30 min in the absence of stabilizing MS2 RNA and over 2 hours in its presence. **(c)** Full LDS-PAGE fluorescent gels representing signals from a pulse-chase experiment using covalent HaloTag fluorescent dyes. U2OS cells that were transiently transfected with plasmids expressing 2xHaloTag-HA-dMCP and MS2 Tornado RNA were stained initially with a JF552 pulse signal (top) before being chased with JF669 (second from top). After gel imaging, protein was transferred to a nitrocellulose membrane, which was stained with anti-HA to view total dMCP levels, displayed in the third image. A second band can be seen to accumulate in all three of these images, which likely represents a decay intermediate that retains both HaloTag and a HA tag. At the bottom, a blot for GAPDH loading control is shown. **(d)** Plotted GAPDH-normalized intensity of JF552 pulse-stained dMCP over time (also normalized to initial concentration) shows that the half life of MS2-stabilized dMCP is approximately 10 hours in this experiment.

#### Supplementary Figure 5

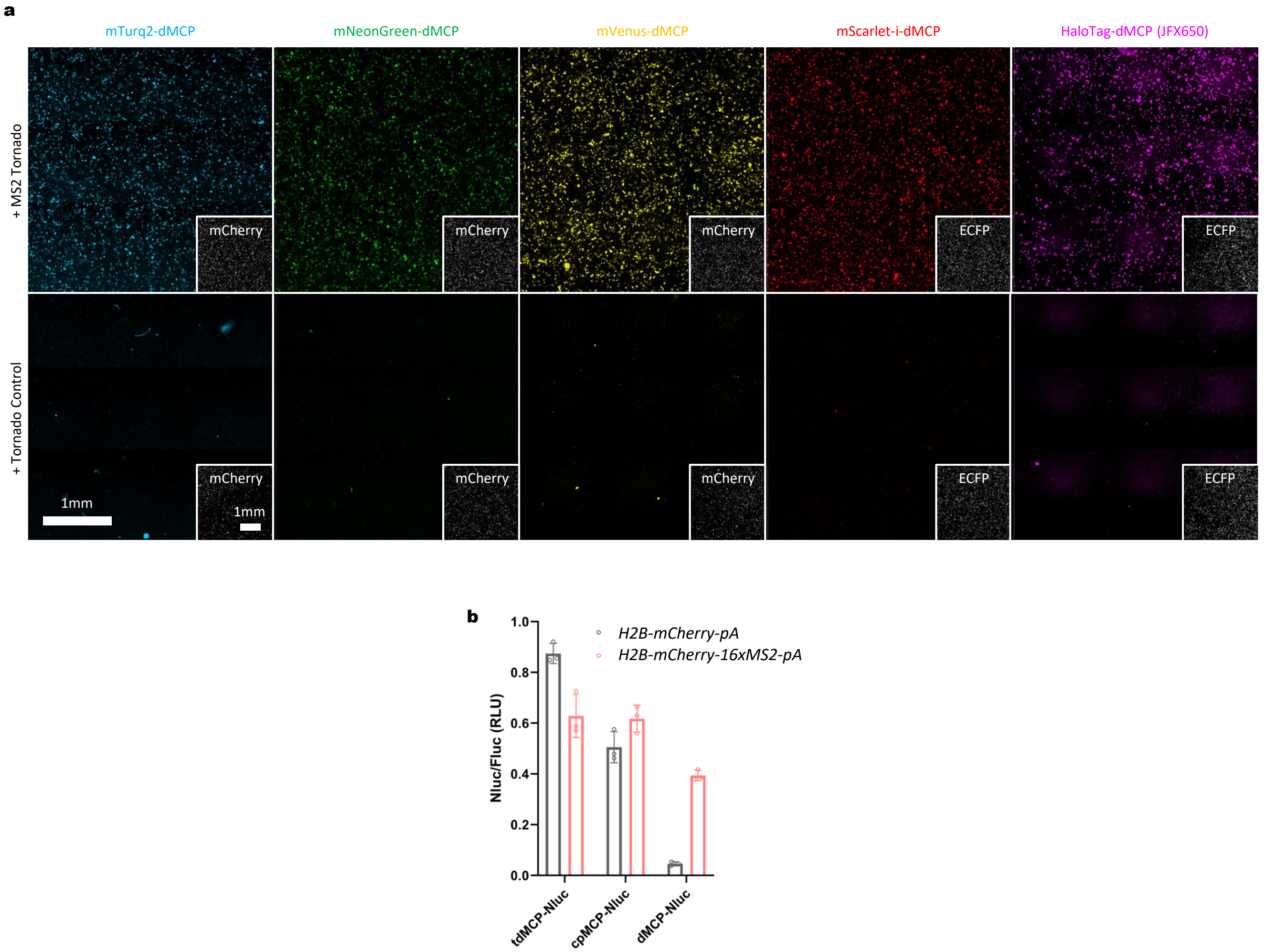

**Supplementary Figure 5 - Extended data of dMCP with various fluorescent and bioluminescent fusion partners.** **(a)** Full-well views of the HEK293FT cells shown in Fig. 1f display dMCP's sustained performance across a large population of transfected cells. Tiled fluorescence microscopy images of five dMCP fusion proteins are shown, with fluorescence emissions ranging from cyan to far-red wavelengths. The signal from mCherry or mTurq2 cotransfection markers are shown in the bottom right windows. **(b)** MS2-RNA stabilization of an NLuc-dMCP fusion construct. Relative NLuc fluorescence was normalized to Fluc co-transfection marker and quantified by plate reader. HEK293FT cells were co-transfected with NLuc-MCP variants and either *H2B-mCherry-pA* (gray circles) or *H2B-mCherry-16xMS2-pA* (red circles). Individual points, bars, and error bars represent the measured intensity, mean and S.D. of n=3 independent transfections.

### Supplementary Figure 6

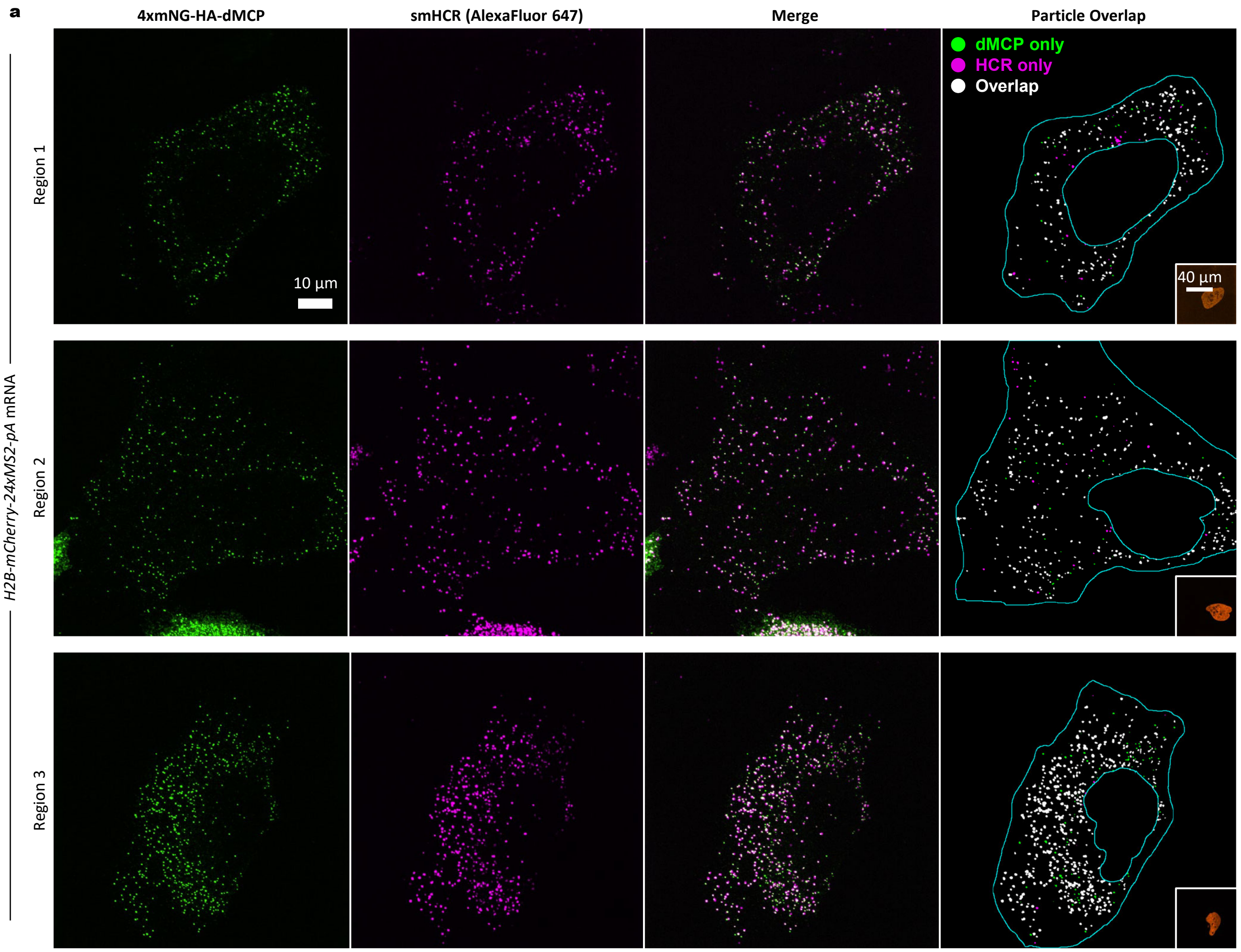


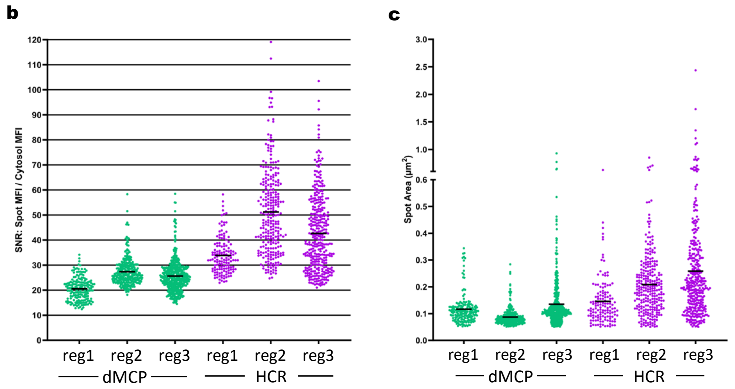


**d**

| Particle type: | reg1 | reg2 | reg3 | sum |
|---|---|---|---|---|
| dMCP colocalizing with HCR particles | 187 | 297 | 396 | 880 |
| dMCP total | 221 | 334 | 480 | 1035 |
| HCR colocalizing with dMCP particles | 166 | 250 | 373 | 789 |
| HCR total | 197 | 277 | 383 | 857 |
| total colocalizing particles | 353 | 547 | 769 | 1646 |
| total particles | 418 | 611 | 863 | 1892 |

| fraction (percent): | reg1 | reg2 | reg3 | avg ± SD |
|---|---|---|---|---|
| dMCP percent colocalization with HCR | 84.6 | 88.9 | 82.5 | 85.0 ± 2.7 |
| HCR percent colocalization with dMCP | 84.3 | 90.3 | 97.4 | 92.1 ± 5.4 |
| total percent colocalization | 84.4 | 89.5 | 89.1 | 87.0 ± 2.3 |

**Supplementary Figure 6 - Extended data comparing dMCP and HCR single-molecule RNA puncta. (a-c)** Maximum intensity projection confocal microscopy images of fixed U2OS cells expressing H2B-mCherry-24xMS2 RNA tagged by 4xmNeonGreen-HA-dMCP and HCR probes targeting mCherry. dMCP signal is shown on the leftmost column (green), HCR signal shown left-of-center (magenta), merged images are shown right-of-center, and an overlay with digitally-labeled cytosolic spots as determined by ImageJ's particle analysis plug-in is shown on the rightmost column. In the overlayed images, green spots are dMCP spots that do not overlap HCR, magenta spots are HCR spots that do not overlap dMCP, and white spots are spots from both channels which overlap. H2B-mCherry signal is shown in the bottom right. **(d)** Signal-to-noise ratio (SNR) of dMCP spots compared with HCR spots across three cells. SNR is measured as the average intensity of a spot divided by the average intensity of the entire cytosolic region excluding spots. Both values are calculated with the average background intensity of the culture medium subtracted. Bars represent the mean of each population. **(c)** Area in square micrometers of dMCP spots in comparison to HCR spots. Bars represent the mean of each population. **(d)** Tabulated values for co-localization between dMCP and HCR spots. dMCP spot counts per cell are a; n=182, b: n=299, c: n=447. HCR spot counts per cell are a: n=138, b: n=269, c: n=345. The lower HCR spot count is primarily due to closely spaced, large puncta that were collectively counted as single foci because they could not be individually resolved.

#### Supplementary FIgure 7

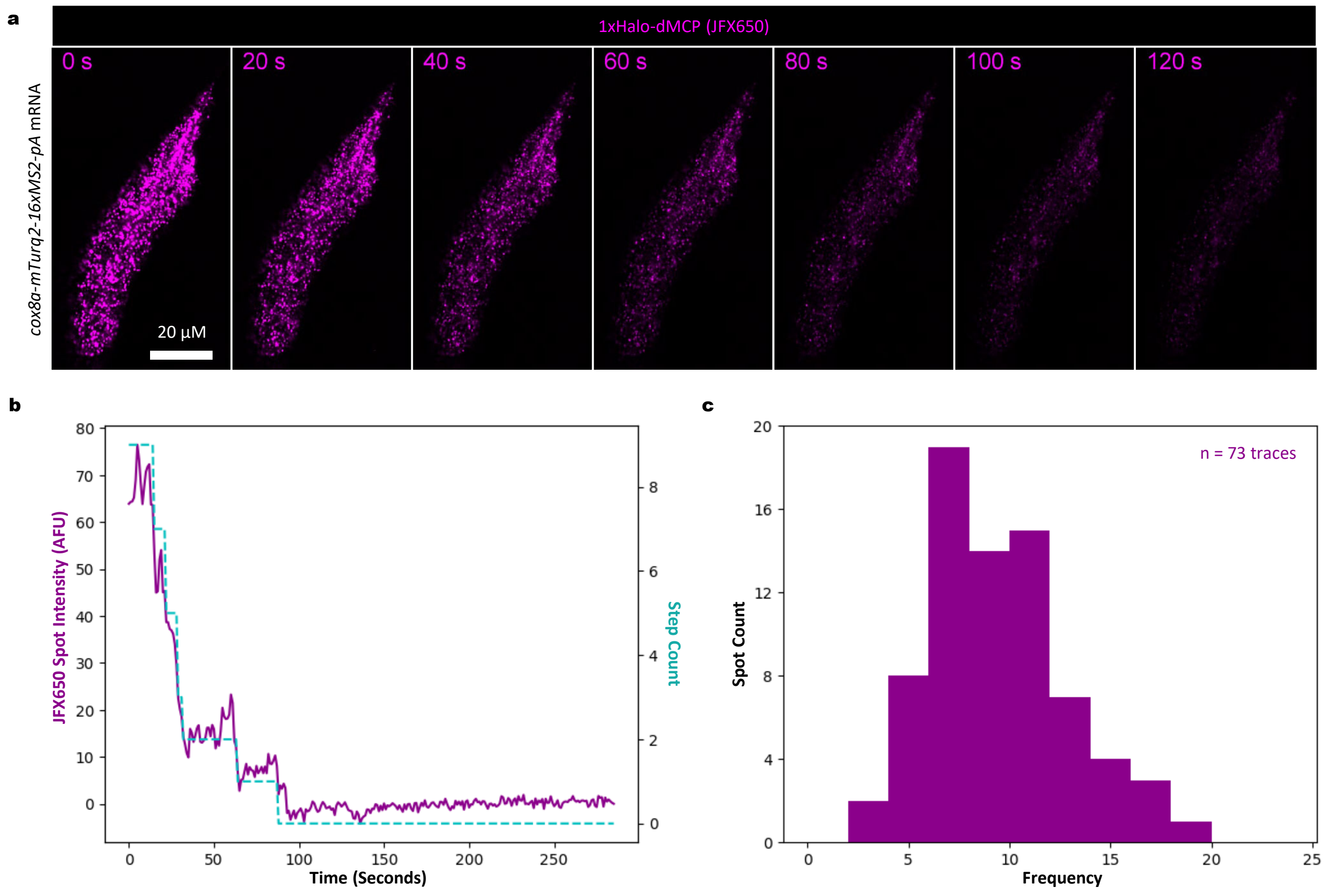

**Supplementary Figure 7 - Step photobleaching analysis of dMCP-tagged mRNA. (a)** Frames from a video of a U2OS cell expressing 1xHaloTag-HA-dMCP tagging *cox8a-16xMS2-pA* mRNA and stained with JFX650-HaloLigand undergoing photobleaching by constant illumination at maximum laser intensity from spinning-disk confocal microscopy. This footage highlights the durability of the JFX650 dye, which allows for long-term imaging of dMCP on far-red channels. **(b)** Example analysis of a punctum undergoing stochastic photobleaching of individual JFX650 dye molecules. Left axis and purple line show an example fluorescent decay trace of a dMCP spot measured with quickPBSA. Right axis and cyan dashed line show a quickPBSA-generated step fit used to predict the quantity of JFX650 dye molecules contained within the spot. **(c)** Histogram showing the distribution of step counts obtained by quickPBSA for spots within the cell. QuickPBSA does not include detected spots which do not have sufficiently clear decay steps in their traces. The data suggests that most spots contain between 8 and 10 dye molecules based on n=73 traces.

### Supplementary Figure 8

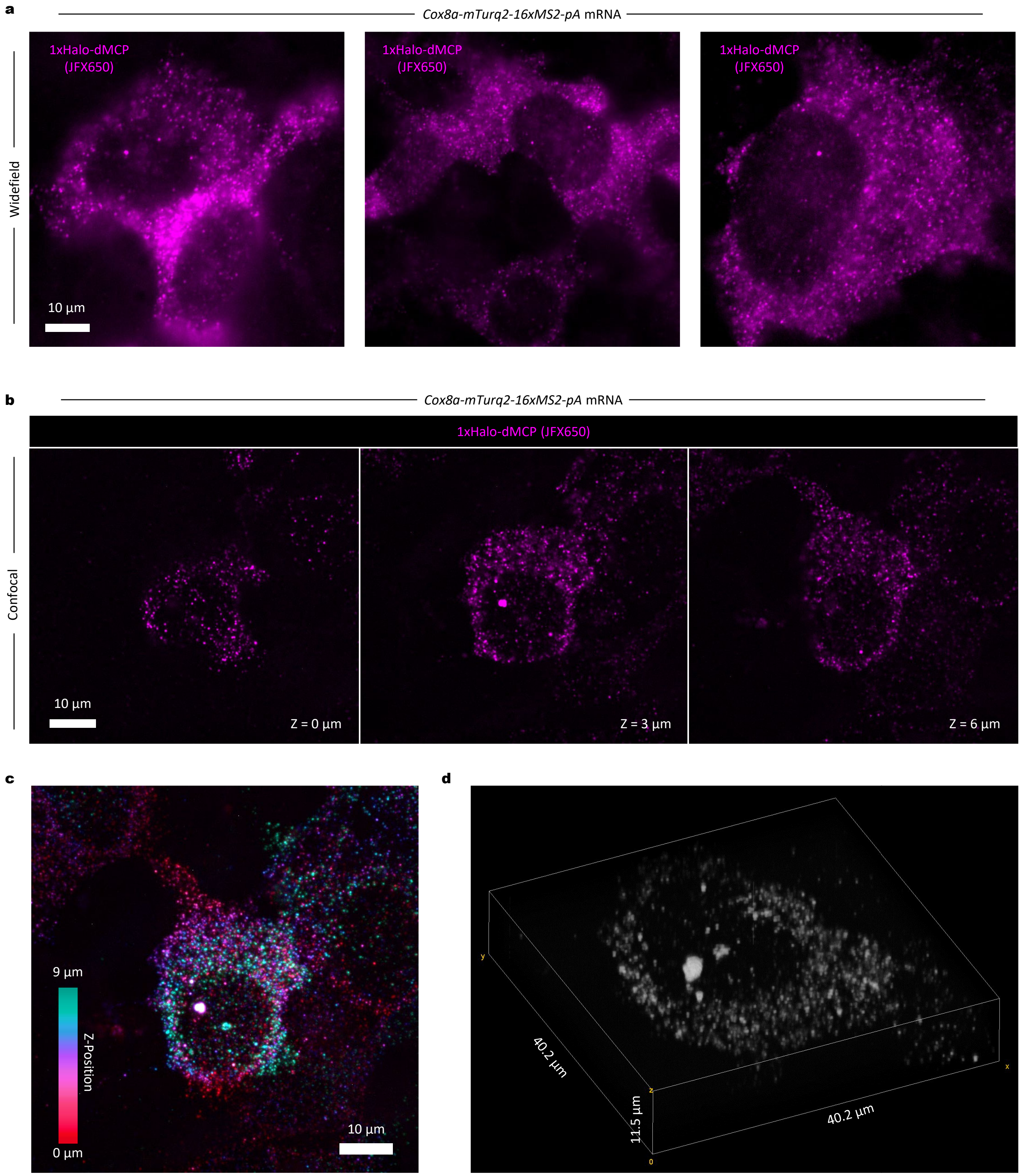

**Supplementary Figure 8 - Nuclear HaloTag-dMCP signal that may correspond to transcriptional burst sites. (a)** Widefield images of HEK293FT cells that were co-transfected to express 1xHaloTag-HA-dMCP and a *Cox8-mTurq2-16xMS2-pA* transcript. Halo-dMCP was stained with JFX650 ligand and cells were fixed before imaging. In a subset of cells, we observed bright clusters of dMCP signal in the nucleus, which may indicate MS2-tagged mRNA within transcriptional burst sites. **(b)** Images of a cell from the same experiment taken as a Z-stack with spinning-disk confocal microscopy. **(c)** A Z-stack projection with dMCP dots color-coded by depth. **(d)** dMCP signal from the same cell rendered in three dimensions.

#### Supplementary Figure 9

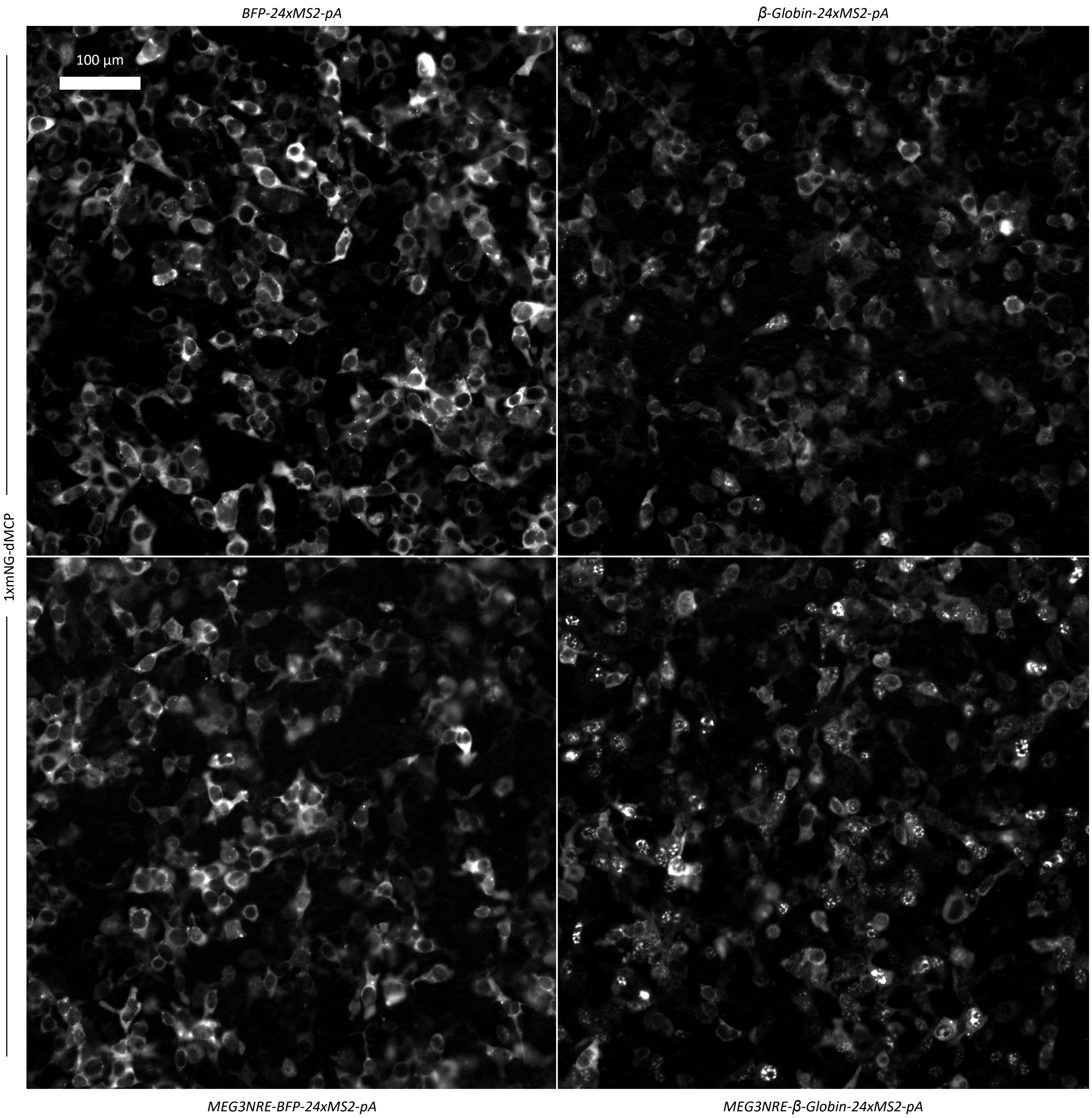

**Supplementary Figure 9 - Expanded images of mRNAs exhibiting varying degrees of nuclear retention as revealed by dMCP.** Here, expanded views of the samples in **Fig 3a.** are shown in order to display a larger population of cells with dMCP-labeled mRNA. HEK293FT were transfected to express mNeonGreen-HA-dMCP and mRNAs that show differing levels of nuclear retention depending on the presence of a MEG3 Nuclear Retention Element and/or introns.

### Supplementary Figure 10

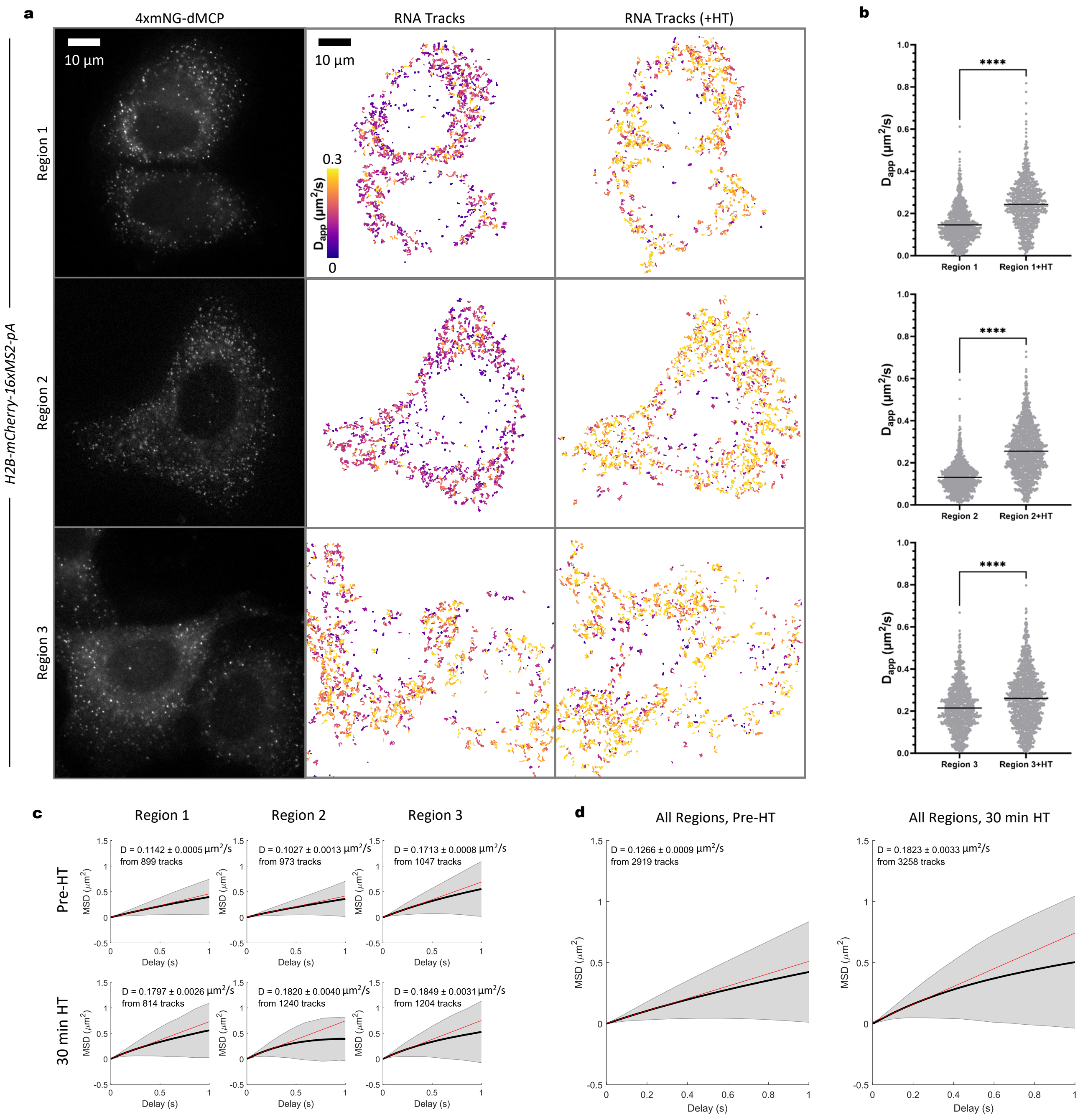

**Supplementary Figure 10 - Extended Data for dMCP tracking of single *H2B-mCherry-16xMS2-pA* transcripts in live cells. (a)** Data from three different live cells expressing 4xmNeonGreen-dMCP and *H2B-mCherry-16xMS2-pA* RNA. Data is displayed in the same format as figure 4: Single frames from videos are displayed in the first column. Color graded overlays of RNA particle traces in translationally active cells, each trace lasting at least 3 frames (0.26 seconds at 0.085 sec/frame), are indicated in the second column. Traces of RNA in the same cells after 30 minutes of translational repression by harringtonine (HT) are in the third column. Color corresponds to apparent diffusion coefficient (calculated using MSD at a time delay of 1 frame) of the trace. **(b)** Apparent diffusion coefficients of particles within each cell before and after harringtonine treatment. All significances determined by unpaired two-tailed Student's t-test: ****P<0.0001. Trace counts: Region 1 n=899, Region 1 HT n=814, Region 2 n=973, Region 2 HT n=1240, Region 3 n=1047, Region 3 HT n=1204. **(c)** MSD curves calculated from above traces using the MSDanalyzer Matlab script. Diffusion coefficients were calculated based on linear fit to the MSD curve and are displayed above. **(d)** MSD curves calculated from the combined traces of all three regions. Diffusion coefficients were calculated based on linear fit to the MSD curve and are displayed above. Trace counts: Pre-HT n=2919, 30 min HT n=3258.

### Supplementary Figure 11

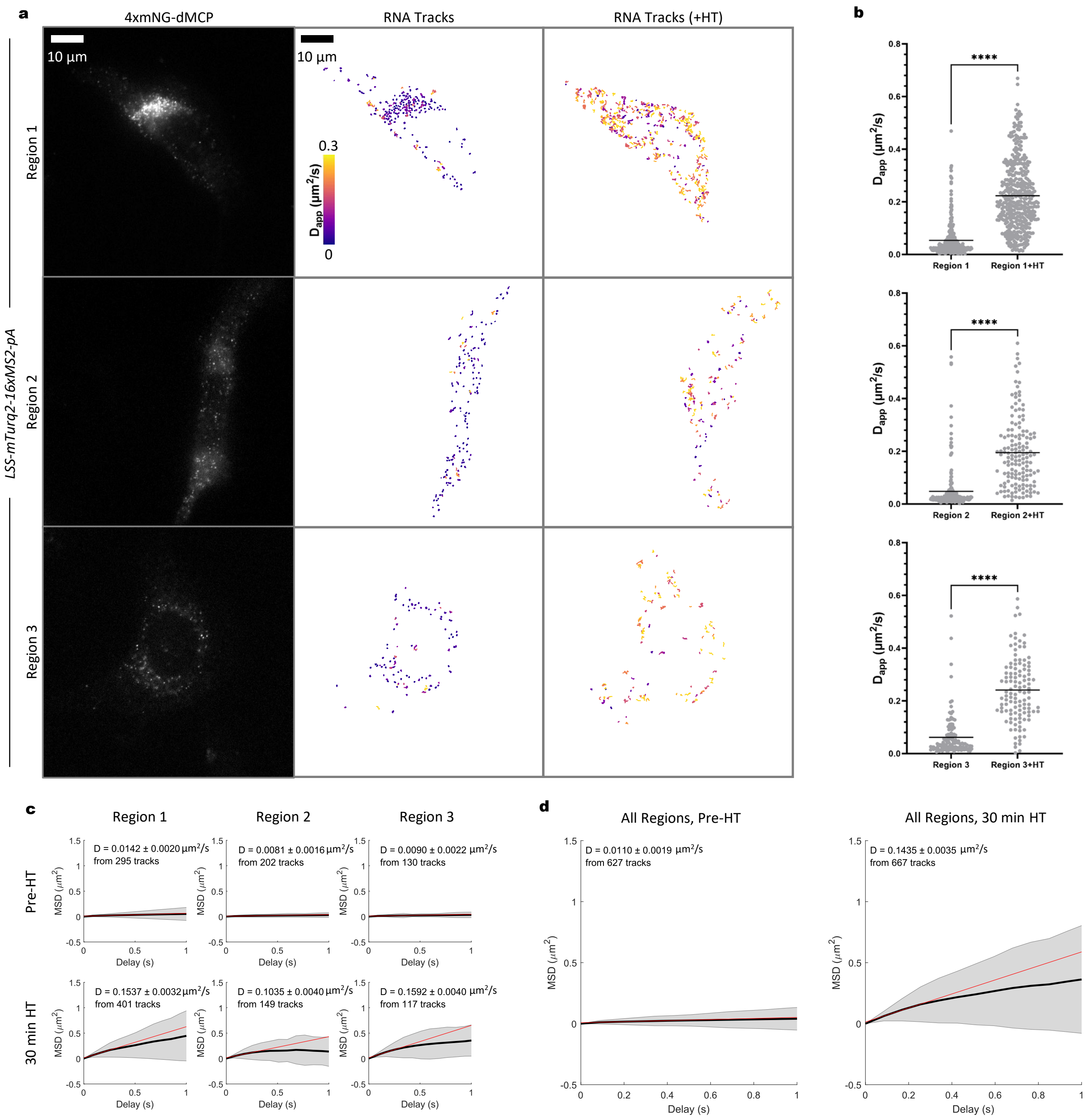

**Supplementary Figure 11 - Extended Data for dMCP tracking of single *LSS-mTurq2-16xMS2-pA* transcripts in live cells.** **(a)** Data from three different live cells expressing 4xmNeonGreen-dMCP and *LSS-mTurq2-16xMS2-pA* RNA. Data is displayed in the same format as figure 4: Single frames from videos are displayed in the first column. Color graded overlays of RNA particle traces in translationally active cells, each trace lasting at least 3 frames (0.26 seconds at 0.085 sec/frame), are indicated in the second column. Traces of RNA in the same cells after 30 minutes of translational repression by harringtonine (HT) are in the third column. Color corresponds to apparent diffusion coefficient (calculated using MSD at a time delay of 1 frame) of the trace. **(b)** Apparent diffusion coefficients of particles within each cell before and after harringtonine treatment. All significances determined by unpaired two-tailed Student's t-test: ****P<0.0001. Trace counts: Region 1 n=295, Region 1 HT n=401, Region 2 n=202, Region 2 HT n=149, Region 3 n=130, Region 3 HT n=117. **(c)** MSD curves calculated from above traces using the MSDanalyzer Matlab script. Diffusion coefficients were calculated based on linear fit to the MSD curve and are displayed above. **(d)** MSD curves calculated from the combined traces of all three regions. Diffusion coefficients were calculated based on linear fit to the MSD curve and are displayed above. Trace counts: Pre-HT n=627, 30 min HT n=667.

### Supplementary Figure 12

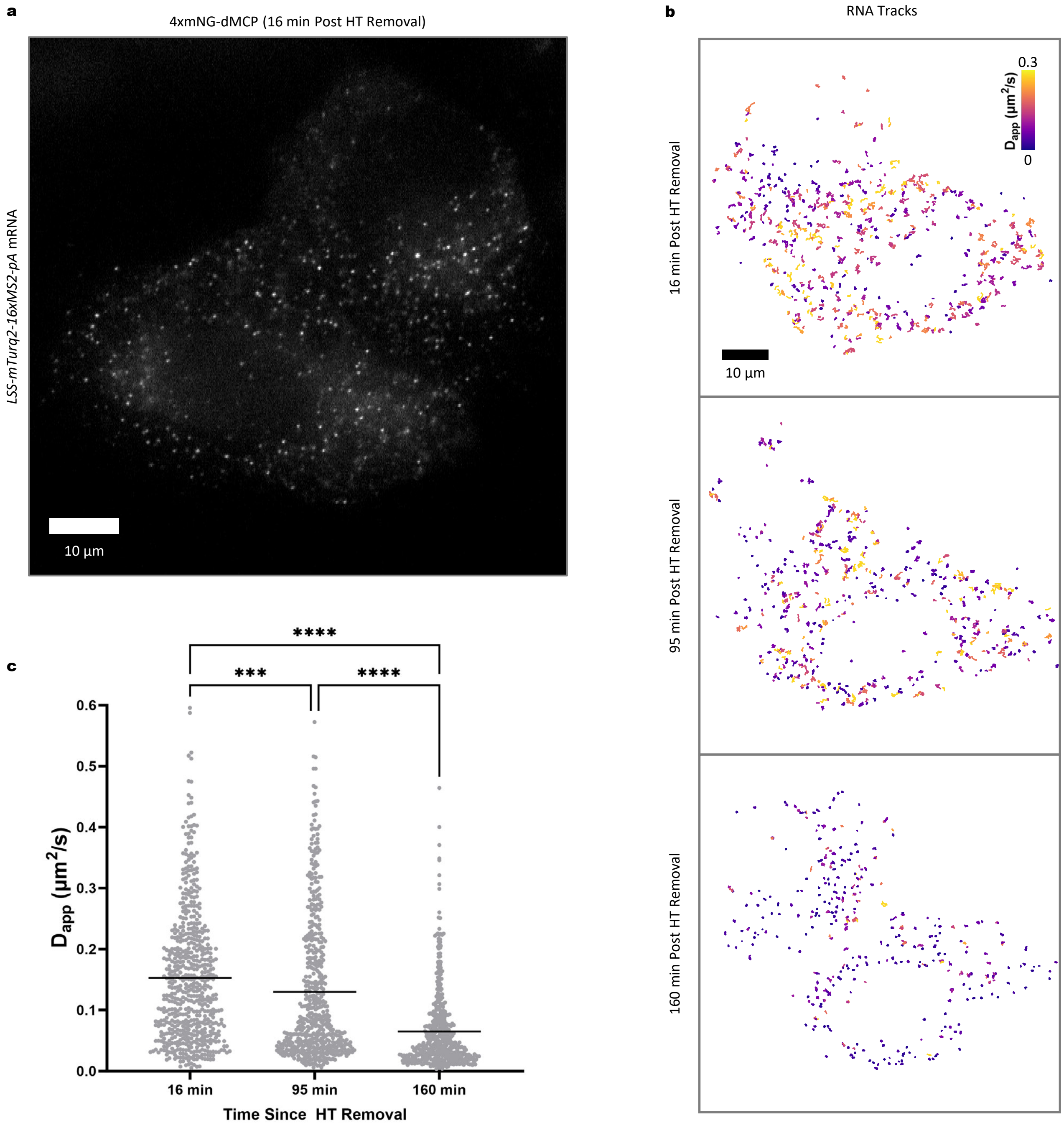

**Supplementary Figure 12 - dMCP imaging to observe the recovery of translation-dependent ER-anchoring in *LSS-mTurq2-16xMS2-pA* transcripts after harringtonine washout. (a)** Image of single RNA molecules shown by dMCP signal in a U2OS cell expressing *LSS-mTurq2-16xMS2-pA* and 4xmNeonGreen-dMCP. Cells were treated with 0.4μM harringtonine for 30 min, followed by washout. The image is from a video taken 16 minutes after washout. **(b)** Color graded overlays of RNA particle traces at several time points after harringtonine washout, each trace lasting at least 3 frames (0.26 seconds at 0.085 sec/frame). Color corresponds to apparent diffusion coefficient (calculated using MSD at a time delay of 1 frame) of the trace. **(c)** The apparent diffusion coefficients of particles within the cell are plotted at each timepoint, showing the recovery of translation-dependent anchoring in *LSS-mTurq2-16xMS2-pA* RNA over several hours. Trace counts: 16 min n=354, 95 min n=299, 160 min n=372. Comparisons are $P_{16vs120min}$ = 0.0394, $P_{95vs120min}$ = 0.0006, and $P_{16vs120min}$ < 0.0001, calculated by ordinary one-way ANOVA

### Supplementary Figure 13

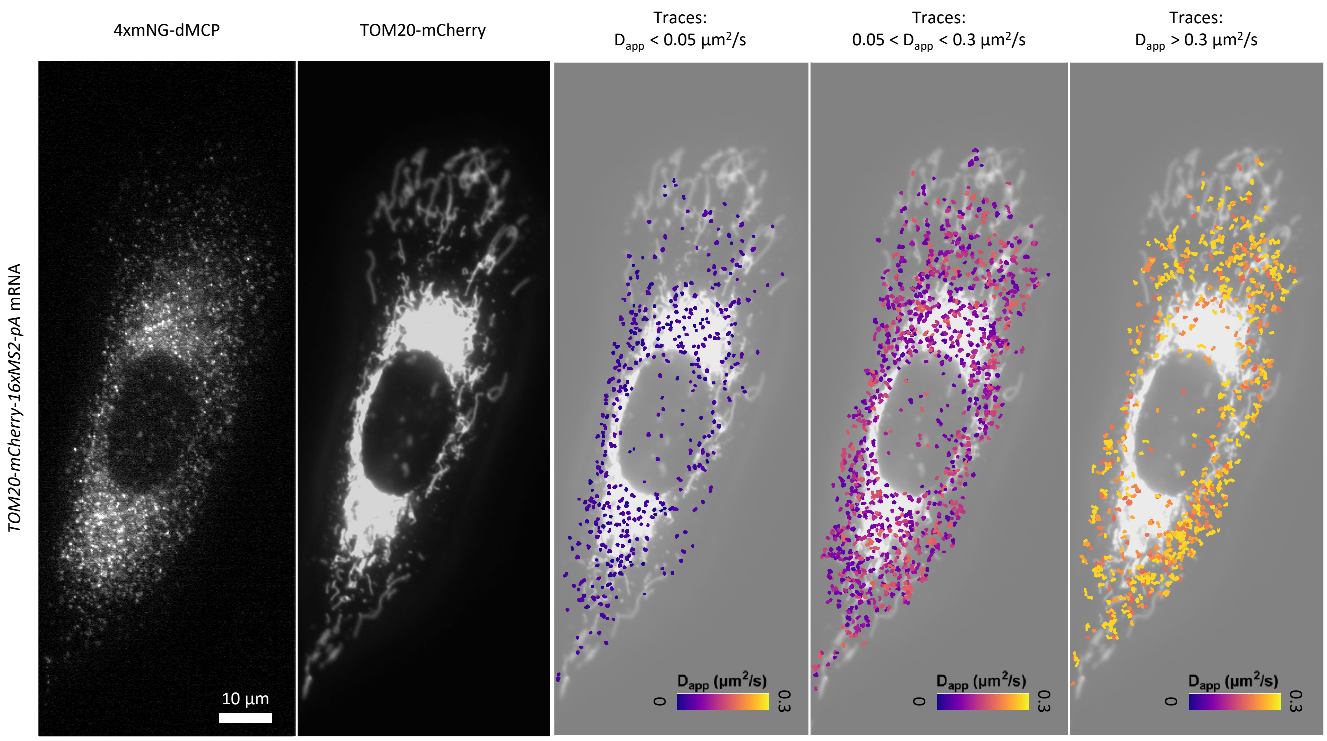

**Supplementary Figure 13 - Subcellular distribution of *TOM20-mCherry-16xMS2* mRNA movements relative to distribution of TOM20-mCherry protein expression.** In the leftmost column, a frame from a recording of dMCP signal from a U2OS cell expressing 4xmNeonGreen-dMCP and *TOM20-mCherry-16xMS2-pA* RNA is shown. Next, an image of TOM20-mCherry expression in the same cell. Following these are overlays of TOM20 signal with dMCP traces color coded by apparent diffusion coefficient as previously described. Three images are presented, with traces exhibiting slow, medium, and fast apparent diffusion rates overlaid respectively. When comparing the overlays, it can be seen that slow traces are more common in subcellular regions with high TOM20-mCherry signal, while fast traces are more common in regions where this signal is reduced.

### Supplementary Figure 14

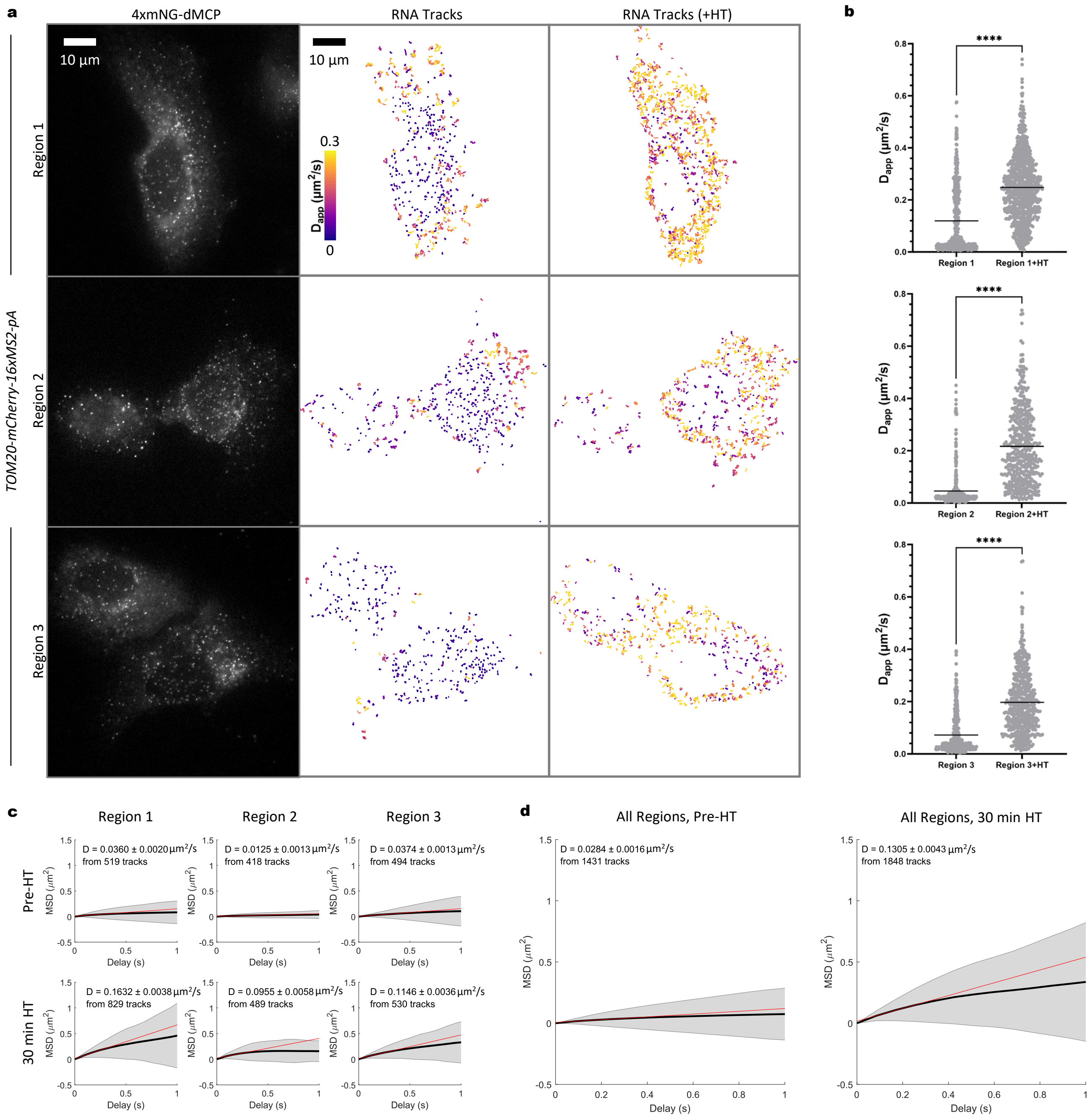

**Supplementary Figure 14 - Extended Data for dMCP tracking of single *TOM20-mCherry-16xMS2-pA* transcripts in live cells. (a)** Data from three different live cells expressing 4xmNeonGreen-dMCP and *TOM20-mCherry-16xMS2-pA* RNA. Data is displayed in the same format as figure 4: Single frames from videos are displayed in the first column. Color graded overlays of RNA particle traces in translationally active cells, each trace lasting at least 3 frames (0.26 seconds at 0.085 sec/frame), are indicated in the second column. Traces of RNA in the same cells after 30 minutes of translational repression by harringtonine (HT) are in the third column. Color corresponds to apparent diffusion coefficient (calculated using MSD at a time delay of 1 frame) of the trace. **(b)** Apparent diffusion coefficients of particles within each cell before and after harringtonine treatment. All significances determined by unpaired two-tailed Student's t-test: ****P<0.0001. Trace counts: Region 1 n=519, Region 1 HT n=829, Region 2 n=418, Region 2 HT n=489, Region 3 n=494, Region 3 HT n=530. **(c)** MSD curves calculated from above traces using the MSDanalyzer Matlab script. Diffusion coefficients were calculated based on linear fit to the MSD curve and are displayed above. **(d)** MSD curves calculated from the combined traces of all three regions. Diffusion coefficients were calculated based on linear fit to the MSD curve and are displayed above. Trace counts: Pre-HT n=1431, 30 min HT n=1848.

### Supplementary Figure 15

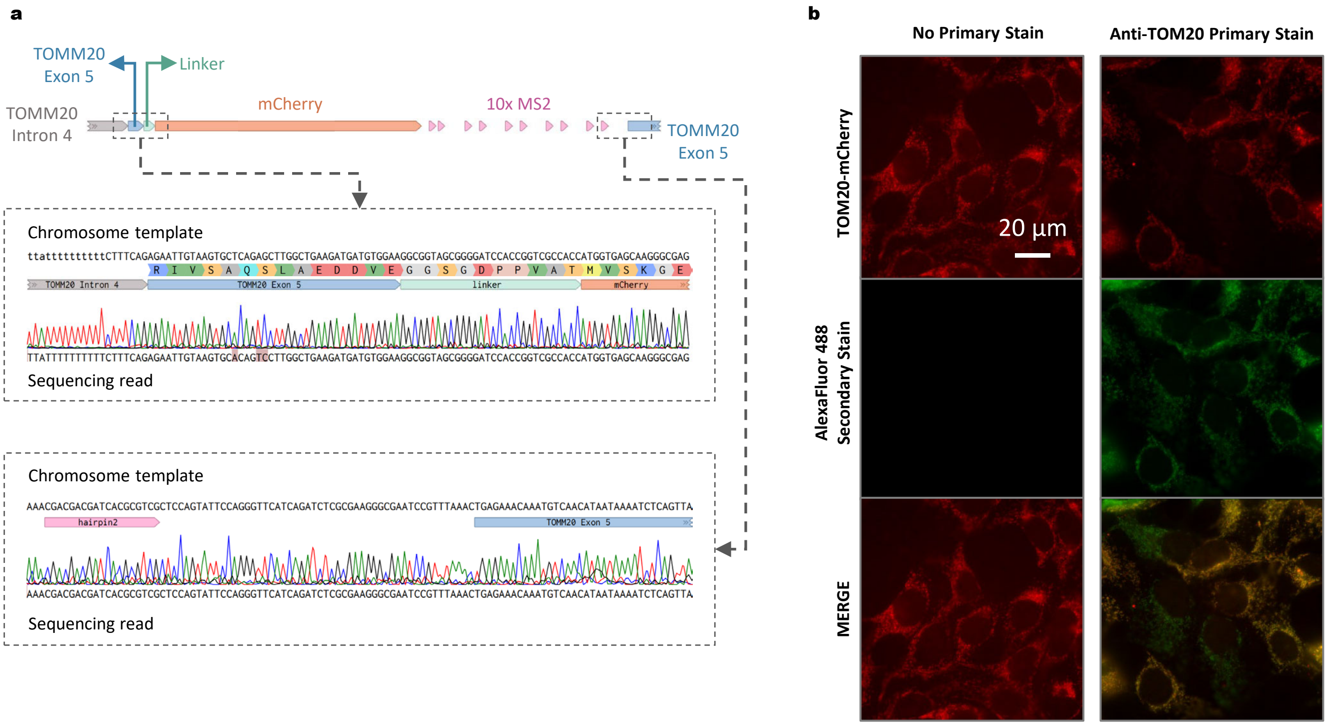

**Supplementary Figure 15– Sequencing and immunostaining confirmation of TOM20-mCherry-10xMS2 CRISPR Knock In. (a)** *top* – gene map of the endogenous TOMM20 locus to where the knock-in of mCherry-10xMS2 cassette was targeted. *Bottom* – Sanger sequencing results of PCR-amplified genomic DNA confirm the successful integration of the MS2 cassette. Silent mutations were introduced to ensure Cas9 targeting was limited to the endogenous locus substrate, and not the post-knock-in product. **(b)** Images of HEK293FT cells with successful CRISPR knock in of mCherry-10xMS2 at the C-terminus of endogenous TOM20. Co-localization of mCherry and TOM20 antibody immunostaining further confirm successful integration and expression of *TOM20-mCherry-10xMS2* and correct targeting of its TOM20-mCherry translation product.

### Supplementary Figure 16

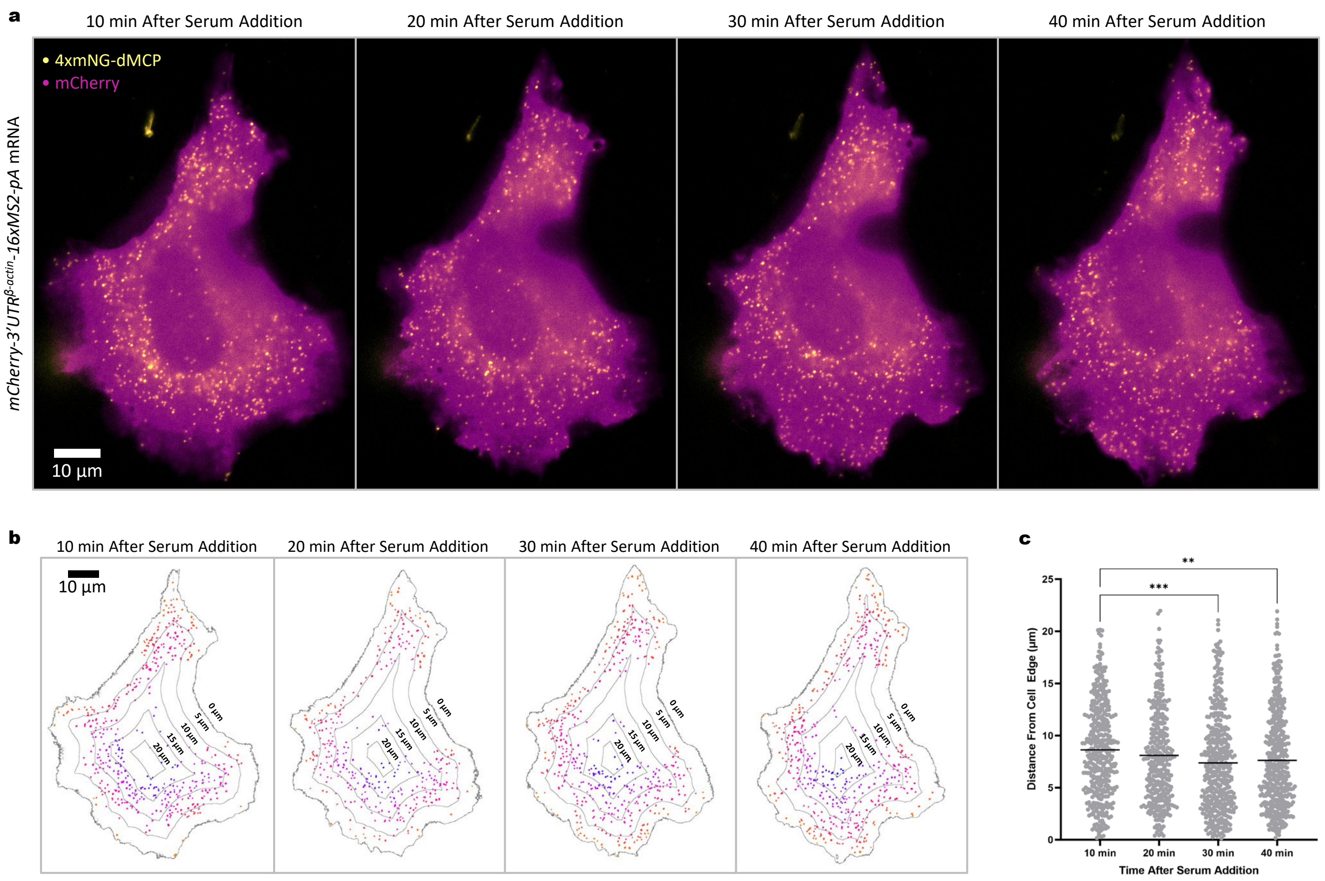

**Supplementary Figure 16 - Additional analysis for β-actin mRNA polarization induced by serum stimulation and imaged within a live cell using dMCP. (a)** Merged images depicting the changing positions of *mCherry-3'UTR*$^{β\text{-}actin}$*-16xMS2-pA* mRNA particles in response to FBS addition to a previously serum-starved U2OS cell. Directed motion towards the cell edge (indicated by mCherry, magenta) by single transcripts labeled by 4xmNG-HA-dMCP (yellow) can be observed. **(b)** Digital render of *mCherry-3'UTR*$^{β\text{-}actin}$*-16xMS2-pA* mRNA particle positions relative to cell boundaries. Particle positions and relative distance from cell-edge was computed using a custom ImageJ plug-in (see extended methods). **(c)** Graph of distance from the cell edge for all particles detected at each time point. Particle counts: 10 min n=404, 20 min n=345, 30 min n=418, 40 min n=431. Comparisons are calculated by ordinary one-way ANOVA, ** P = 0.0073, *** P = 0.0005.

#### Supplementary Figure 17

a

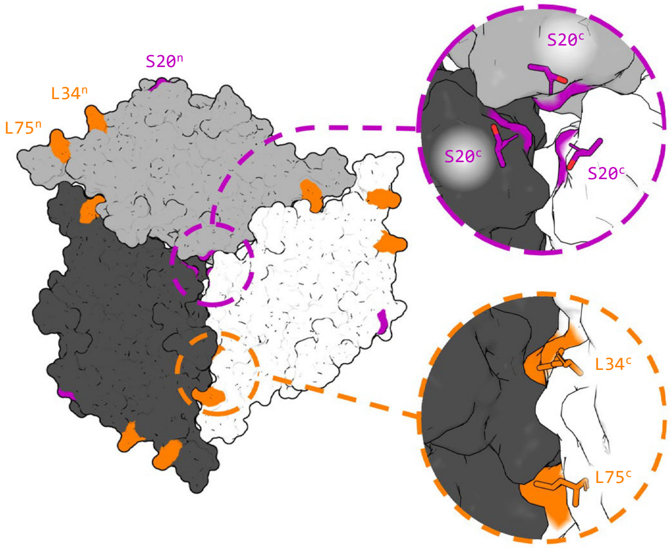


b

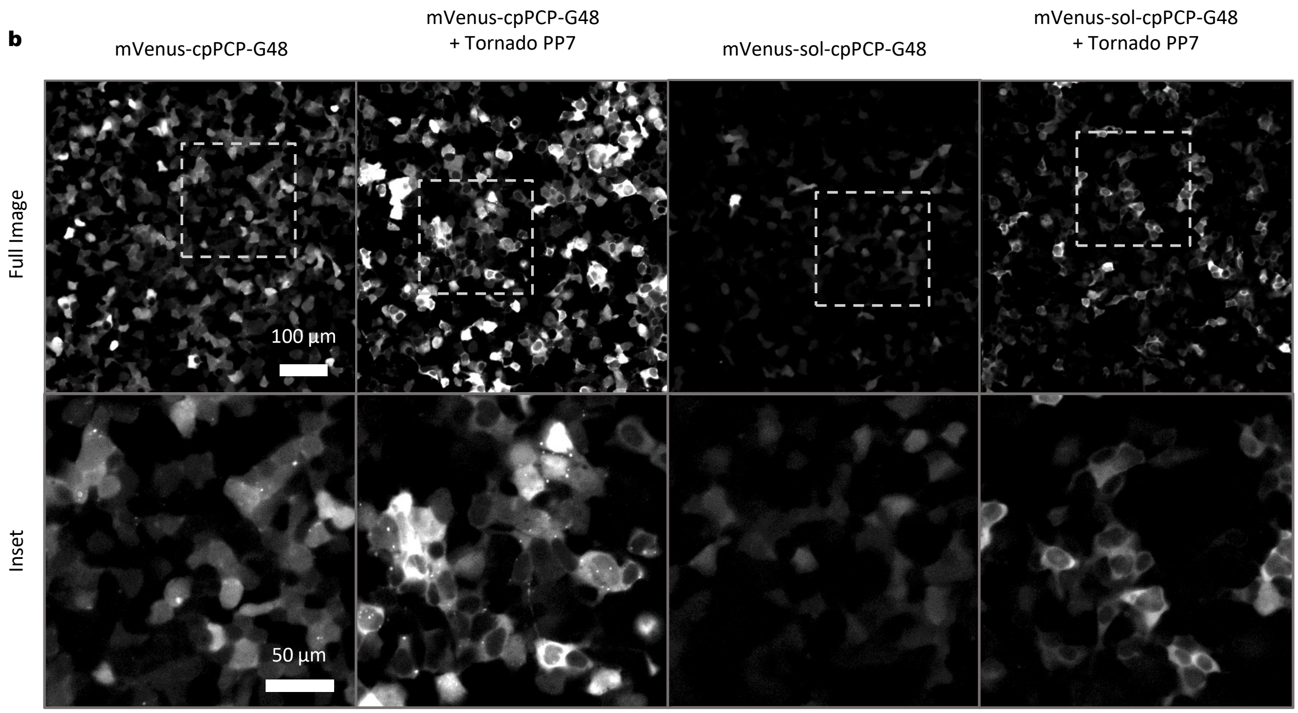

**Supplementary Figure 17 - Development of solubilized circularly-permuted PP7 coat proteins by targeted mutagenesis. (a)** Protein structure of a 3-way interface formed by FG-loop containing tandem dimeric PP7 coat proteins (PDB – 6N4V). Residues are labeled with n or c superscript to denote whether they correspond to the N-terminal or C-terminal monomeric halves of the fused dimer. Leucine residues that appear to be involved with nonpolar interactions are highlighted in orange, and these residues are mutated to alanine in the solubilized version of cpPCP-G48. Serine residues at the core of the trimeric interaction are highlighted in purple, and these residues are mutated to arginine in sol-cpPCP-G48 to interfere with trimeric protein packing. **(b)** Images of mVenus-cpPCP-G48 and mVenus-sol-cpPCP-G48, expressed with or without PP7 tornado RNA. MS2 RNA-dependent nuclear exclusion indicates binding of the tdPCP circular-permuted variants to PP7 RNA. Punctate aggregates can be seen for mVenus-cpPCP-G48, but the same aggregates are not visible for sol-cpPCP-G48.

#### Supplementary Figure 18

a

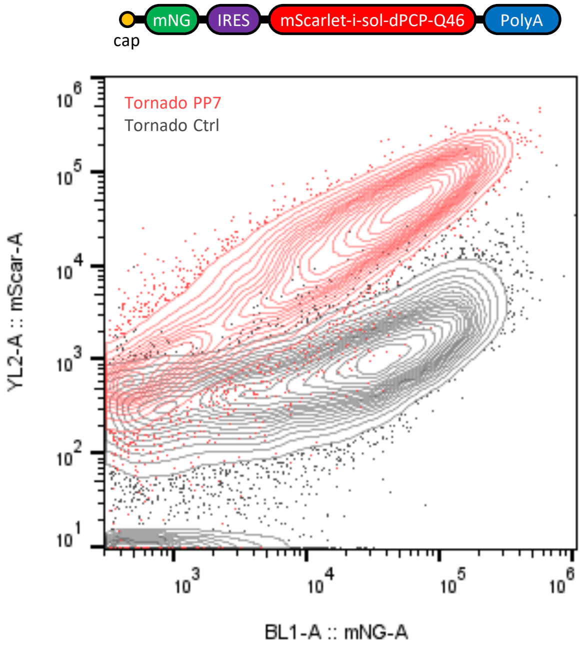

**Supplementary Figure 18 - Flow cytometry data of the optimized sol-dPCP-Q46 variant.** *Top* - Schematic of the reporter construct, with cap-dependent translation of a stable mNeonGreen expression reporter and IRES-dependent translation of the optimal sol-dPCP-Q46 (‘dPCP’) fused to mScarlet. *Below* - Representative flow cytometry traces from HEK293FT cells expressing the above reporter and PP7 Tornado (red) or Tornado Control (gray) RNA.

#### MATERIALS AND METHODS

##### General cloning

New DNA constructs were generated using standard cloning procedures via either T4 DNA Ligase or Gibson Assembly reactions. Restriction enzymes, ligases, and assembly mixtures were obtained from New England Biolabs (NEB). Plasmids containing repeating DNA sequences (including those with MS2 or PP7 repeat arrays), plasmids encoding the described circular RNAs, and lentiviral transfer vectors were transformed and prepped using a recombination-deficient NEB Stable *E. coli* strain (NEB) as the transformation host. Transformed NEB Stable cells were grown on agar plates or in liquid cultures at 30° C. Plasmids were confirmed via Sanger sequencing before use.

##### Cloning of circular RNA encoding plasmids

Plasmids encoding the circular tornado-MS2, tornado-PP7, and tornado-control RNA sequences were generated by modification of pAV-U6+27-Tornado-F30-TAR Variant-1 (AddGene Plasmid #129406). Briefly, the parent vector was digested at *NheI* and *KpnI* sites and dephosphorylated using Quick CIP (NEB) before isolation of the cleaved backbone by DNA gel extraction. Inserts based on annealed oligonucleotide pairs (Integrated DNA Technologies, IDT) were phosphorylated using T4 Polynucleotide Kinase before ligation into the cleaved plasmid backbone using T4 DNA Ligase. The resulting plasmids contain the following indicated DNA sequences inserted between the *NheI* and *KpnI* sites of the original tornado-encoding backbone (lowercase letters correspond to the indicated hairpin-forming sequences):

| | |
|---|---|
| tornado-MS2: | 5'-AAAgcacgagcatcagccgtgcCGA-3' |
| tornado-PP7: | 5'-AAAggagcagacgatatggcgtcgctccCGA-3' |
| tornado-control: | 5'-ATTAGCTCCGAGCCCGA-3' |

##### Mammalian cells

Mammalian cells were maintained in a humidified incubator at 37° C with 5% $CO_2$. HEK293-FT (ThermoFisher, R70007) and U2OS (Sigma-Aldrich, 92022711-1VL) cells were grown in media based on high-glucose (4.5 g/L) Dulbecco's Modified Eagle Medium (DMEM) containing sodium pyruvate (Cytiva SH30285.01) supplemented with 10% (v/v) fetal bovine serum (FBS, typically either Cytiva Characterized Fetal Bovine Serum, Canadian Origin (Cytiva, SH30396.03), or Corning Regular Fetal Bovine Serum (Corning, 35010CV)), along with nonessential amino acids (LifeTechnologies), Glutamax (LifeTechnologies), and penicillin-streptomycin (ThermoFisher).

Primary neonatal human dermal fibroblast cells (HDF-Neo, Lonza CC-2509, Lot# 0000670357) were grown in Fibroblast Growth Medium-2 containing complete supplements associated with the FGM-2 BulletKit (Lonza, CC-3132). Early passage HDF-neo cells used (passage 5 or earlier).

##### DNA transfections

Transfections were performed using the Lipofectamine 3000 reagent (ThermoFisher). Transfection complexes were prepared according to the supplier's protocol. Amounts of DNA and levels of co-transfected plasmids were varied based on individual experimental applications as described in the sections below.

##### MS2-binding validation using NLS-cpMCP-S52 and initial test of dMCP-S52 degradation

HEK293-FT cells were grown on fibronectin-coated (FN-coated) 18-well chambered coverglass slips (Cellvis, C18SB-1.5H). Coating was performed using 10 µg/ml FN in PBS for 1 h at room temperature, followed by rinsing three times with PBS before adding cells. HEK293-FT cells were seeded into the glass plates at 50,000 cells per well and transfected with 100 ng of vector encoding mVenus-NLS-HA-cpMCP-S52, mVenus-HA-cpMCP-S52, or mVenus-HA-dMCP-S52 driven by a minCMV promoter, along with 100 ng of a vector encoding *H2B-mCherry-pA* or *H2B-mCherry-16xMS2-pA* transcripts driven via CMV promoters. Cells were transfected in suspension by combining transfection complexes with cells at the time of cell seeding. Cells were imaged by widefield microscopy at two days post-transfection.

**Flow cytometry**

Flow cytometry analyses were performed using an Attune-NxT Flow Cytometer (ThermoFisher). Transfected cells were analyzed at ~48 hours post-transfection. For cells grown in 96 well plates, cell suspensions were prepared by aspiration of the growth medium followed by incubation in 50 µL of EDTA-containing 0.25% Trypsin solution (ThermoFisher). Trypsinization proceeded at 37°C for no longer than 5 min, after which the reaction was quenched by the addition of 200 µL of complete growth medium. The resulting cell suspension was used in flow cytometry analyses. Detection voltages were adjusted on an experiment-by-experiment basis. A representative example of the implemented gating procedures is provided in **Supplementary Fig. 2**. Live cells were identified by FSC-A and SSC-A gating. Singlet cells were identified via gating based on FSC-A versus FSC-H. Transfection-positive cells were defined as those exhibiting mNeonGreen and BFP emission at intensities above that of the top 99.9% of non-transfected control cells. FlowJo™ v10 Software was used for all flow cytometry analyses.

**Optimization of degron-positionings**

Transfected HEK293-FT cells were used in the dMCP and dPCP optimization screens. Cells were seeded into TC-treated 96-well plates at 80,000 cells per well. Levels of mNG, mScarlet-i, and BFP were quantified by flow cytometry at two days post-transfection.

For MCP-based coat proteins, cells were transfected with plasmid mixtures containing pcDNA3-based bicistronic constructs encoding a CMV-driven cap-dependent mNeonGreen (mNG) in combination with encephalomyocarditis virus (EMCV) IRES driven mScarlet-i-fused coat protein sequences. Cells were co-transfected with 10 ng of bicistronic plasmid in combination with 100 ng of plasmid encoding either *BFP-24xMS2-pA* or *BFP-pA*, both driven from CMV promoters.

For PCP-based coat proteins, a CMV promoter-driven mNG was co-expressed in combination with mScarlet-i-fused coat protein sequences, also via an EMCV-IRES and the pcDNA3 backbone. Cells were co-transfected with DNA mixtures containing three plasmids, including: 10 ng of bicistronic plasmid, 100 ng of plasmid encoding either tornado-PP7 or tornado-control driven from U6 promoters, and 100 ng of a BFP-encoding plasmid as a cotransfection marker.

**Analysis of fluorescent protein (FP)-dMCP fusions**

HEK293-FT cells were grown on FN-coated 8-well chambered coverglass slips (Cellvis, C8-1.5H-N). Cells were transfected in suspension by combining 200,000 cells per well with Lipofectamine 3000 transfection complexes. For imaging FP-dMCP fusions stabilized by tornado-MS2 (as shown in **Fig. 1f** and **Supplementary Fig. 4a**), transfection mixtures were prepared by mixing 200 ng of plasmid encoding a minimal CMV promoter (minCMV)-driven reporter-dMCP fusion, 200 ng of plasmid encoding a U6 promoter-driven tornado construct (tornado-control or tornado-MS2, as indicated in the figures), and 50 ng of plasmid encoding a fluorescent cotransfection marker expressed from a UBC promoter (mCherry or ECFP, selected based on spectral compatibility and as indicated in the figures). Cells were imaged at two days post-transfection under live conditions in FluoroBrite imaging media (A1896702, ThermoFisher) containing the same supplements as cell growth media.

**Proteasome inhibition**

HEK293-FT cells were transfected with bicistronic constructs encoding mNG in combination with IRES-regulated mScarlet-dMCP. Transfections were performed in 96-well plates using 100,000 cells per well. The day after transfection, cells were treated with MG-132 (Selleckchem) or lactacystin (Santa Cruz Biotechnology), each using 10 µM treatment concentrations. Proteasome inhibition proceeded by incubation with each drug for 6 hours prior to analyzing mNG and mScarlet levels using flow cytometry as described above.

**Half-life analyses using cycloheximide**

HEK293-FT cells were transfected with 30 ng of plasmid encoding a UBC promoter-driven mNG-HA-dMCP in combination with 300 ng of plasmid encoding U6 promoter-driven tornado-MS2 or tornado-control. DNA mixtures were combined with 270 ng of salmon sperm DNA as filler prior to preparing transfection complexes using Lipofectamine 3000. Transfections were performed using cell suspensions, with cells seeded into 48-well TC-treated plates at 250,000 cells per well. CHX pulse treatments were performed at one day post-transfection by treatment with CHX at a dose of 15 µM for 0, 30, 60, and 120 minute durations. Cells were lysed using 1X NuPAGE LDS sample buffer (ThermoFisher) and lysates were stored frozen until analysis by western blotting (see below).

**Fluorescent pulse-chase analyses**

U2OS cells were seeded into a 48-well plate at 50,000 cells per well and co-transfected with 100 ng plasmid encoding 2xHaloTag-HA-dMCP (driven from UBC promoter) in combination with 100 ng of a plasmid encoding tornado-MS2 or tornado-control (driven from U6 promoters). Pulse-chase labeling was initiated at 2 days post-transfection by exchange of cells into growth media containing 200 nM of a JF552-based HaloTag ligand. Staining with the JF552-based ligand proceeded for 1 h at 37° C, after which cells were washed 3x5 min with fresh growth media. Chase labeling was then initiated by exchanging cells into growth media containing 200 nM of a JF669-based HaloTag ligand. Cells were grown in the JF669-containing solution for the durations indicated in figures; samples were collected by lysing cells in 1xLDS sample buffer. Cell lysates were separated via SDS-PAGE as described in the associated section below. Fluorescence emissions from the covalently stained proteins were recorded via direct in-gel detection using an iBright Imaging System (ThermoFisher). Note that JF552- and JF669-based ligands were selected on the basis of their spectral compatibility and also due to their non-fluorogenic nature (permitting the detection of denatured protein conjugates in SDS-PAGE gels). To confirm protein loading and to acquire loading control densities for normalizing fluorescent band intensities, proteins were transferred to nitrocellulose membranes after fluorescence detection. Blocked membranes were then probed with antibodies against 2xHalo-HA-dMCP (using anti-HA, for detection of total 2xHalo-HA-dMCP levels) and GAPDH (using anti-GAPDH-HRP). Details for blotting procedures and the utilized antibodies are provided in the subsequent section.

**SDS-PAGE and western blotting**

Cell lysates were prepared by aspiration of growth media from individual culture wells followed by rinsing cells once with PBS before direct lysis in 1X NuPAGE LDS Sample Buffer (ThermoFisher).

Lysates were sonicated to shear genomic DNA and reduce sample viscosity, then stored frozen at -20° C until use. Sample aliquots were reduced by adding NuPAGE Sample Reducing Agent (ThermoFisher) and denatured by heating at 70° C for 5-10 minutes. Reduced and denatured samples were separated by SDS-PAGE. Transfer to nitrocellulose membranes was carried out using an iBlot2 transfer device (ThermoFisher). Membranes were blocked using a blocking buffer based on PBS containing 0.05% Tween-20 (volume/volume) (PBS-T) containing nonfat dry milk, dissolved at 5% weight/volume.

Blocked membranes were probed with a mouse monoclonal anti-HA antibody (Clone 6E2, Cell Signaling Technologies) using a 1:1,000 volumetric dilution in blocking buffer. Probing with anti-HA proceeded overnight at 4° C, with rocking. Membranes were washed three times with PBS-T (for 5 min per wash) prior to probing with a horse anti-mouse secondary HRP-conjugate (Cell Signaling Technologies, 4047) in PBS-T at a dilution of 1:3,000. Probing with the secondary antibody proceeded for 1 h in blocking buffer at room temperature prior washing again three times with PBS-T. Signals from HRP conjugates were developed using freshly prepared SuperSignal West Pico PLUS Chemiluminescent Substrate mixtures (ThermoFisher), with chemiluminescent recording using an iBright Imaging System (ThermoFisher). Probed membranes were stripped using Pierce Restore Western Blot Stripping Buffer (ThermoFisher) and re-blocked with blocking buffer prior to detecting loading control proteins, via either a rat anti-GAPDH HRP conjugate (BioLegend, 607903) using 1:10,000 dilution in PBS-T, or a mouse anti-β-actin-HRP conjugate (BioLegend 643808) at a 1:3,000 dilution, also in PBS-T. Probing with loading control antibodies proceeded for 1 hour at room temperature before washing and chemiluminescent development as described. Band intensities were quantified using ImageJ.

**Preparation of cells for single-molecule RNA imaging under live or fixed conditions**

U2OS cells were seeded into FN-coated glass-bottom 8-well imaging dishes at 50,000 cells per well. Cells were co-transfected with 100 ng of plasmid encoding the indicated dMCP-fusion proteins driven from UBC promoters in combination with 100 ng of plasmid encoding the indicated MS2-tagged transcripts, also expressed from UBC promoters. Cells were imaged two days post-transfection. HDF-neo cells were prepared for single-molecule imaging using the lentiviral production and transduction procedures described below, passaged twice, then seeded on FN-coated glass-bottom 8-well dishes at 50,000 cells per well in media containing 200 ng/µL doxycycline (Sigma-Aldrich). HDF-Neo cells were imaged after two days under doxycycline induction.

**Imaging of MS2-tagged mRNAs via 4xmNeonGreen-dMCP and smHCR**

U2OS cells were prepared as described above. At two days post-transfection, cells were rinsed once with PBS prior to fixation using a pre-warmed 4% paraformaldehyde (PFA) solution. Working solutions of the fixative were prepared by diluting methanol-free PFA from 16% (w/v) solutions stocks (ThermoFisher, 28906) into PBS. Fixation proceeded for 20 minutes at 37° C before removing the fixative and rinsing once with TBS-glycine buffer to quench residual PFA. Cells were rinsed an additional two times with PBS following the quenching step. Fluorescence *in situ* hybridization (FISH) was performed using an smHCR protocol based on that described by Choi *et al.* (*Development,* 145(12), dev165753)[26]. Briefly, antisense probes against *mCherry* mRNA (Molecular Instruments) were used in combination with AlexaFluor-647 (AF647)-conjugated “B1” HCR amplifier hairpins (Molecular Instruments). Hybridization and amplification steps were performed as described by Choi *et al.*, with modifications: (1) cells were permeabilized using Triton X-100 (0.2% v/v in PBS) in place of ethanol-based permeabilization to minimize signal loss from the expressed fluorescent proteins, and (2) final imaging was done using cells immersed in

PBS, in place of antifade mounting solution. HCR-labeled cells were imaged using a spinning disk confocal microscope (described below).

**Confocal imaging of dMCP- and HCR-labeled transcripts**

Confocal images were taken on an Andor Dragonfly 505 spinning disk confocal microscope equipped with a Zyla 4.2 plus sCMOS Camera with a 2x zoom lens and a 100x/1.45 numerical aperture oil objective lens. Signals from 4xmNeonGreen-dMCP were detected through a Chroma ET525/50m emission filter using 488 nm laser excitation at 1000 ms excitation time. Signals from H2B-mCherry were detected through a Chroma ET620/60x emission filter using 561 nm laser excitation at 1000 ms excitation time. Signals from AF647-conjugated HCR hairpins were detected through a Semrock FF01-698/70 emission filter using 641 nm laser excitation at 200 ms excitation (timed shorter to limit the photobleaching of AF647 chromophores). Z-stack recordings were carried out using a 200 nm step size; maximum intensity projections involved combining between 10-30 individual Z-slices, depending on sample dimensions.

**Image analysis for SNR, particle area, and particle colocalization**

Z-stack images from the Dragonfly Microscope were converted to maximum intensity projections for ImageJ analysis of SNR, particle area, and particle colocalization. The 'Analyze Particles' function was used to identify dMCP and HCR spots, their mean intensities, and their areas. The cytosolic intensity was measured as the mean intensity of pixels within the cytosol, excluding pixels in the nucleus and pixels in spots. Background intensity was measured as the mean intensity of pixels in a large rectangular region of unoccupied space outside the cell. Signal-to-noise was calculated as the difference between spot intensity and background intensity divided by the difference between cytosolic intensity and background intensity. Spot colocalization was calculated as the portion of dMCP spots whose perimeter was within a one-pixel radius of the HCR spot perimeter and vice versa.

**Widefield imaging**

Widefield images were taken with a Zeiss AxioObserver Z1 microscope equipped with an HXP 120V halogen lamp as the excitation source. Images were recorded using a Prime95B sCMOS camera (Teledyne Photometrics) and the ZEN imaging software (Black Edition, Zeiss). Images were recorded through a 10x/0.3-NA air objective lens, a 20x/0.8-NA air objective lens, or a 63x/1.4-NA oil objective lens. mTurquoise2 and ECFP were visualized under 'ECFP' filter cube settings (Chroma 49001; ET436/20x, T455lp, ET480/40m); mNeonGreen and mVenus under 'EYFP' settings (Chroma 49003; ET500/20x, T515lp, ET535/30m); mCherry, mScarlet-i, JF570, and JF585 under 'mCherry' settings (Chroma 49008; ET560/40x, T585lpxr, ET630/75m); JFX646 and JFX650 under 'narrow-excitation Cy5' settings (Chroma 49009; ET640/30x, T660lpxr, ET690/50m). Exposure times of 50 ms were used for live imaging single-RNA dynamics with 4xmNG-dMCP, 100 ms exposures were used for live imaging of 2xmNG-dMCP, and 200 ms exposures were used for live imaging of RNA with 2xHaloTag-dMCP stained by JFX650-based ligands, or 4xmNeonGreen-dPCP. For any other imaging, the exposure time that maximized the camera's dynamic range was used, ranging between 2 and 2000 ms depending on the reporter and imaging context.

**Staining and imaging of HaloTag-dMCP fusions**

JaneliaFluor (JF)-containing HaloTag ligands based on chloroalkane-modified dyes were gifts from Dr. Luke Lavis of Janelia Farm. Cells expressing HaloTag-fused proteins were stained under live conditions at 37° C. Cells were stained in growth media containing 200 nM dye. To ensure their full dissolution, dyes were diluted into pre-warmed media aliquots and mixed vigorously by pipetting and brief vortexing prior to application to cells. For imaging under live conditions, cells were stained in dye-containing media for 40 minutes with incubation at 37° C prior to removing the staining solution and rinsing with fresh, pre-warmed media (3x5 min). For cells imaged under fixed conditions, cells were fixed using 4% PFA immediately after removal of cell staining media. PFA fixation and subsequent rinsing was carried out as described above. Stained live cells were imaged live in fully supplemented FluoroBrite-based media. Fixed cells were imaged in PBS.

**Bioluminesence measurements**

For the bioluminescent assay, 100,000 HEK293FT cells were transfected in suspension and plated on a 96-well white flat-bottom TC-treated microwell plate. Transfection mixtures were prepared using the following DNA amounts for each transfected well: 10 ng of plasmid encoding the indicated NLuc-MCP variant driven by minCMV promoters, 50 ng of plasmid encoding CMV-driven H2B-mCherry (either with or without an inserted 16xMS2 array), and 50 ng of a plasmid encoding a CMV-driven Firefly Luciferase (FLuc) as a transfection control. End point luminescence values were obtained two days after transfection using Nano-Glo Dual-Luciferase Reporter (NanoDLR) Assay System (Promega) according to the manufacturer's instructions. Briefly, growth media from individual wells was replaced with 40 µL of Opti-MEM (ThermoFisher) before adding 40 µL of ONE-GLO EX substrate. After incubating at room temperature for 5 min, FLuc luminescence values were measured with a SpectraMax M5 Multimode Microplate Reader (Molecular Devices) using one read area and a one-second integration time per well. NanoDLR Stop & Glo Substrate was diluted 1:100 into Stop & Glo Buffer before adding 40 µL to each well to quench FLuc luminescence and induce NLuc luminescence. After incubating at room temperature for 15 min, NLuc luminescence values were measured using the microplate reader with the same acquisition settings as FLuc. Relative luminescence values were obtained by dividing NLuc luminescent values by the FLuc values for each well.

**Imaging of nuclear transcripts using 1xHaloTag-dMCP**

For imaging nuclear-localized transcripts (as in **Supplementary Fig. 6**), HEK293-FT cells were seeded into FN-coated glass bottom 8-well imaging dishes at 100,000 cells per well. Cells were transfected with 200 ng of plasmid encoding 1xHaloTag-dMCP expressed from a minCMV promoter in combination with 200 ng of plasmid encoding *Cox8-mTurq2-16xMS2-pA*, also via a minCMV promoter. The next day, cells were stained with JFX650 HaloTag ligand using media containing the dye at 200 nM. Staining proceeded for 40 min, followed by one rinse with PBS before immediately fixing cells using 4% PFA solution. Fixation proceeded as described in the preceding sections. Cells were rinsed with PBS three times before imaging in PBS.

**Step photobleaching analysis**

U2OS cells were seeded into FN-coated glass bottom 8-well imaging dishes at 50,000 cells per well. Cells were transfected in suspension using transfection complexes prepared with 100ng each of plasmid DNA encoding 1xHaloTag-dMCP and *Cox8-mTurq2-16xMS2-pA*, both driven from UBC promoters. Two days after transfection, cells were stained with a JFX650-based HaloTag ligand prior to rinsing with PBS and fixation with 4% PFA, both as described above. JFX650-stained dMCP spots were recorded by spinning disc confocal microscopy using 1000 ms exposure times. Photobleaching proceeded via continuous excitation at maximum laser intensity over a 5 minute duration, with continuous recording of signal. TrackMate, via ImageJ, was used to identify centroids of each spot. The quickPBSA python library was then used to generate photobleaching traces from the trackmate centroid coordinates and the photobleaching videos. QuickPBSA was also used for automated filtering and subsequent analysis of these traces.

**Imaging of nuclearly retained mRNAs**

Images in **Fig. 3a** and **Supplementary Fig. 8** are widefield microscopy images of transfected HEK293-FT. Cells were plated on an FN-coated glass bottom 8-well imaging dish at 200,000 cells per well and co-transfected with plasmid mixtures containing 200 ng of plasmid encoding mNG-HA-dMCP in combination with 200 ng plasmid encoding the indicated transcripts. In these analyses,mNG-dMCP was expressed from a minCMV promoter, with expression of the indicated MS2-tagged transcripts driven by UBC promoters.

Images in **Fig. 3b** and **Fig. 3c** are of U2OS cells, also grown in FN-coated imaging wells, with cells seeded at a density of 50,000 cells per well. Cells were transfected with 100 ng of plasmid encoding 1xHaloTag-dMCP in combination with 100 ng of plasmid encoding *MEG3NRE-β-Globin-24xMS2-pA,* both expressed from UBC promoters. Cells were stained with a JFX650-based HaloTag ligand followed by rinsing, fixation, and permeabilization, as described previously. Cells were imaged in PBS containing DAPI (4′,6-diamidino-2-phenylindole) at 30 nM as a counterstain.

**Lentiviral transduction**

Lentiviral particles were produced using a second-generation vector system. HEK293-FT cells were grown to 90% confluence in a six-well dish and transfected with 750 ng of pLV-based transfer plasmid encoding 2xmNeonGreen-dMCP on a UBC promoter or *CFP-24xMS2-pA* on a doxycycline-inducible Tre3G promoter, as well as 1.25 µg of psPax2 packaging plasmid, and 1.25 µg of pVSVG envelope plasmid. Transfection media was replaced with fresh media the next day. Virus-containing supernatants were harvested 24 h and 48 h later. Supernatants were filtered through a 0.45 µm filter before immediate use in cell transduction or storage at -80° C. Primary neonatal human dermal fibroblast cells were transduced by direct addition of filtered supernatant into cell culture.

**Harringtonine treatment**

For experiments involving single-molecule imaging with pre and post harringtonine (HT; HY-N0862, MedChemExpress) treatment comparisons, U2OS cells were prepared as described above and initially imaged in pre-warmed FluoroBrite media. Cell positions were recorded for repeated viewing at later time points. Next, a solution of 400µM concentrated HT in fluorobrite was pipetted directly into wells, without moving the plate, for a final concentration of 4 µM HT. Cells were then incubated in the scope at 37C for 30 minutes before recording initiation-blocked transcript trajectories at pre-recorded cell positions. For the harringtonine removal experiment, cells were treated with 0.4 µM harringtonine for 30 minutes, then washed three times with fresh media, followed by imaging in FluoroBrite media with intermittent recording over a two-hour timespan.

**Analysis of particle movements**

TrackMate (via ImageJ) was used to detect and analyze particle movements from live-cell dMCP recordings of single RNA molecules. Videos of 4xmNeonGreen-dMCP emissions were recorded for 40 frames at 0.085 sec/frame using a 50 ms exposure time and with pixel width of 109 nm. The videos shown in **Fig. 5f**, taken with 2xHalo-dMCP and 4xmNeonGreen-dPCP, are both 20 frames at 0.2255 sec/frame (200ms exposure) and 218 nm pixel width (these pixels are larger because 2x2 binning was used to increase sensitivity). Single particles were detected using TrackMate's LoG detector for 3-pixel diameter spots and filtered for quality. Tracking was performed with a simple LAP tracker. Maximum particle displacement over one frame for

4xmNeonGreen-dMCP videos was set to 5 pixels (0.55 µm). For 2xHalo-dMCP and 4xmNeonGreen-dPCP, this displacement was set to 3 binned pixels (0.65 µm). Gaps were closed over a maximum of 3 frames and either 5 pixels or 3 binned pixels as described above. Only tracks with at least 3 localizations were included in the analysis. Note that setting minimum track durations at too high a value (above 5) biases track detection towards slower moving particles, as they remain in focus for longer durations compared to faster moving particles (with the latter tending to diffuse out of the visible z-dimension).

Data from trackmate analyses was exported in XML format and the MSDanalyzer matlab package was used to calculate mean-squared displacement values from the exported files. The calculated diffusion coefficients ($D$) shown in the bottom sections of **Supplementary Fig. 10, 11, and 13** were calculated from the combined MSD curves of all traces involved. Apparent diffusion coefficients ($D_{app}$) of single traces were calculated as described in previous works[34], using the mean squared displacement at a time delay of one frame, dimensionality ($n$) of two, and the time delay ($t$) in seconds per single frame, according to the following formula:

$$\mathrm{MSD} = D_{app} \cdot 2nt$$

Individual particle traces were plotted as vector graphics and color coded by apparent diffusion coefficient using an in house Python script.

#### Serum stimulation

U2OS cells were co-transfected with plasmids encoding 4xmNG-dMCP and *mCherry-3'UTR*$^{\beta\text{-}actin}$*-16xMS2-pA* in preparation for live single molecule imaging as described above. The next day, cells were starved of serum by exchange into serum-free growth media (prepared in a similar manner as standard growth media but with omission of FBS). After 24 hours under serum starvation, cells were imaged with visualization of the labeled transcripts and recording of cell locations as 2D coordinates within imaging wells. The serum starved cells were then stimulated by exchange into standard FBS-containing growth media. Transcript locations were then imaged via widefield microscopy every 10 minutes over a one hour period.

The digital analyses seen in **Supplementary Fig. 15b and 15c** were performed using ImageJ. First, images of 4xmNG-labeled RNA and mCherry emissions were treated with a difference of gaussians filter (DoG) to enhance edge detection, then the 'analyze particles' function was used to define cell boundaries using the filtered mCherry image and the boundaries of RNA punctae using the filtered mNG image. The distance of individual 4xmNG punctae from the edge of the cell was calculated using an ImageJ-based macro.

#### Generation of CRISPR-tagged cells

Donor DNA containing homology arms corresponding to the human TOMM20 locus was a gift from the Allen Institute for Cell Science (AICSDP-8:TOMM20-mEGFP, AddGene, Plasmid #87423). The plasmid was modified by replacing the GFP insert with a sequence corresponding to mCherry-10xMS2. The donor was integrated into HEK293-FT cells using guide RNAs targeting the endogenous TOMM20 locus (at annealing sites 5'-AATTGTAAGTGCTCAGAGCT-3' and 5'-TGGTAGTTGAGCAGCTCTGGGGG-3') and on-target genomic integration was confirmed using PCR and Sanger sequencing. Integration resulted in the translation of TOMM20-mCherry protein from 10xMS2-tagged mRNA. The correct localization of TOMM20-mCherry protein was confirmed through immunostaining with a mouse monoclonal anti-TOMM20 antibody (clone F-10; sc-17764, Santa Cruz Biotechnology). For immunostaining, cells were fixed in 4% PFA (as described above) and permeabilized by treatment with PBS containing Triton-X 100 (0.2%, v/v) for 5 min at room temperature. Immunostaining was then performed overnight at 4° C via incubation in Immunofluorescence Blocking Buffer (12411S, Cell Signaling Technology) containing diluted primary antibody at a 1:1,000 dilution. Primary-stained samples were then washed three times in PBS-T (5 min per rinse) before secondary detection was carried out using an AlexFluor488-conjugated anti-mouse secondary antibody (A-11001, ThermoFisher) at 1:2000 dilution. Secondary antibody staining proceeded at room temperature for one hour in PBS-T, before rinsing three times in PBS-T (5 min per rinse) followed by imaging in PBS-T.

**Two-color imaging using dMCP and dPCP**

The fluorescence images displayed in **Fig. 5d and 5e** are of co-transfected HEK293-FT cells. Cells were plated on an FN-coated glass bottom 8-well imaging dish at 200,000 cells per well and co-transfected with plasmid mixtures encoding the indicated proteins and RNAs. Plasmids encoding minCMV-driven mScarlet-dMCP and mNG-dPCP were transfected at 100 ng each; plasmids encoding RNAs based on U6-driven circular RNAs (tornado MS2, tornado PP7, and tornado control) or the CMV-driven *MEG3NRE-β-Globin-24xMS2-pA* transcript, were also transfected at 100 ng each.

The images shown in Fig. 5f are of a U2OS cell in a FN-coated imaging well that was seeded at 50,000 cells per well and co-transfected with plasmid mixtures encoding the indicated proteins and RNAs. The transfection mixture contained 100ng each of four plasmids: one encoding 2xHalo-dMCP, one encoding 4xmNG-dPCP, one encoding *H2B-mCherry-24xMS2*, and one encoding *LSS-mTurq2-18xPP7*, all driven by UBC promoters.

**CAPTION FOR SUPPLEMENTARY MOVIES 1-9**

**Supplementary Video 1 -** U2OS cells transfected to express 4xmNeonGreen-dMCP and *H2B-mCherry-16xMS2-pA*. Cells are imaged before and after 30 minute treatment with the translational repressor harringtonine. 50ms exposure time.

**Supplementary Video 2 -** HDF-Neo primary cells transduced to express 2xmNG-dMCP and *CFP-24xMS2-pA* mRNA. 100ms exposure time.

**Supplementary Video 3 -** U2OS cell transfected to express 4xmNeonGreen-dMCP and *LSS-mTurq2-16xMS2-pA*. In this cell, dMCP-tagged *LSS-mTurq2-16xMS2-pA* mRNA appears largely anchored to regions of high mTurq2 expression. 50ms exposure time for both channels.

**Supplementary Video 4 -** U2OS cell transfected to express 4xmNeonGreen-dMCP and *LSS-mTurq2-16xMS2-pA*. Cells are imaged before and after 30 minute treatment with the translational repressor harringtonine. 50ms exposure time.

**Supplementary Video 5 -** U2OS cell transfected to express 4xmNeonGreen-dMCP and *TOM20-mCherry-16xMS2-pA*. In this cell, dMCP-tagged *TOM20-mCherry-16xMS2-pA* mRNA moves more slowly in subcellular regions of high TOM20 expression. 50ms exposure time for both channels.

**Supplementary Video 6 -** U2OS cell transfected to express 4xmNeonGreen-dMCP and *TOM20-mCherry-16xMS2-pA*. Cells are imaged before and after 30 minute treatment with the translational repressor harringtonine. 50ms exposure time.

**Supplementary Video 7 -** HEK293FT cell with CRISPR knock-in of *TOM20-mCherry-10xMS2* at the endogenous TOM20 locus. Arrows indicate regions of TOM20-mCherry signal colocalizing with dMCP-labeled TOM20 transcripts over several seconds. The two videos were recorded simultaneously, with 500ms exposure time used for imaging 4xmNG-dMCP and 100ms exposure time used for imaging TOM20-mCherry.

**Supplementary Video 8 -** U2OS cell transfected to express 4xmNeonGreen-dMCP and *H2B-mCherry-18xPP7-pA*. 200ms exposure time.

**Supplementary Video 9 -** U2OS cell transfected to express 2xHalo-dMCP, 4xmNeonGreen-dPCP, *H2B-mCherry-24xMS2-pA*, and *LSS-mTurq2-18xPP7-pA*. 200ms exposure time for both channels. Videos were recorded back-to-back to avoid loss of framerate that occurs due to switching of filter cubes.

**AMINO ACID SEQUENCES FOR THE REPORTED PROTEIN DOMAINS**

| **CONSTRUCT** | **AMINO ACID SEQUENCE** |
|---|---|
| tdMCP | ASNFTQFVLVDNGGTGDVTVAPSNFANGIAEWISSNSRSQAYKVTCSVRQSSAQNRKYTIKVEVPKGAWRSYLNMELTIPIFATNSDCELIVKAMQGLLKDGNPIPSAIAANSGIYA*NFTQFVLVDNGGTGDVTVAPSNFANGIAEWISSNSRSQAYKVTCSVRQSSAQNRKYTIKVEVPKGAWRSYLNMELTIPIFATNSDCELIVKAMQGLLKDGNPIPSAIAANSGIY* |
| cpMCP-S52 | AQNRKYTIKVEVPKGAWRSYLNMELTIPIFATNSDCELIVKAMQGLLKDGNPIPSAIAANSGIYA*NFTQFVLVDNGGTGDVTVAPSNFANGIAEWISSNSRSQAYKVTCSVRQSSAQNRKYTIKVEVPKGAWRSYLNMELTIPIFATNSDCELIVKAMQGLLKDGNPIPSAIAANSGIY*ANFTQFVLVDNGGTGDVTVAPSNFANGIAEWISSNSRSQAYKVTCS**VRQSS** |
| dMCP-S52 | AQNRKYTIKVEVPKGAWRSYLNMELTIPIFATNSDCELIVKAMQGLLKDGNPIPSAIAANSGIYA*NFTQFVLVDNGGTGDVTVAPSNFANGIAEWISSNSRSQAYKVTCSVRQSSAQNRKYTIKVEVPKGAWRSYLNMELTIPIFATNSDCELIVKAMQGLLKDGNPIPSAIAANSGIY*ANFTQFVLVDNGGTGDVTVAPSNFANGIAEWISSNSRSQAYKVTCS**VRQSSRRRG** |
| dMCP-S51 | AQNRKYTIKVEVPKGAWRSYLNMELTIPIFATNSDCELIVKAMQGLLKDGNPIPSAIAANSGIYA*NFTQFVLVDNGGTGDVTVAPSNFANGIAEWISSNSRSQAYKVTCSVRQSSAQNRKYTIKVEVPKGAWRSYLNMELTIPIFATNSDCELIVKAMQGLLKDGNPIPSAIAANSGIY*ANFTQFVLVDNGGTGDVTVAPSNFANGIAEWISSNSRSQAYKVTCS**VRQSRRRG** |
| **dMCP-Q50**<br><br>**‘dMCP’** | AQNRKYTIKVEVPKGAWRSYLNMELTIPIFATNSDCELIVKAMQGLLKDGNPIPSAIAANSGIYA*NFTQFVLVDNGGTGDVTVAPSNFANGIAEWISSNSRSQAYKVTCSVRQSSAQNRKYTIKVEVPKGAWRSYLNMELTIPIFATNSDCELIVKAMQGLLKDGNPIPSAIAANSGIY*ANFTQFVLVDNGGTGDVTVAPSNFANGIAEWISSNSRSQAYKVTCS**VRQRRRG** |
| dMCP-R49 | AQNRKYTIKVEVPKGAWRSYLNMELTIPIFATNSDCELIVKAMQGLLKDGNPIPSAIAANSGIYA*NFTQFVLVDNGGTGDVTVAPSNFANGIAEWISSNSRSQAYKVTCSVRQSSAQNRKYTIKVEVPKGAWRSYLNMELTIPIFATNSDCELIVKAMQGLLKDGNPIPSAIAANSGIY*ANFTQFVLVDNGGTGDVTVAPSNFANGIAEWISSNSRSQAYKVTCS**VRRRRG** |
| dMCP-V48 | AQNRKYTIKVEVPKGAWRSYLNMELTIPIFATNSDCELIVKAMQGLLKDGNPIPSAIAANSGIYA*NFTQFVLVDNGGTGDVTVAPSNFANGIAEWISSNSRSQAYKVTCSVRQSSAQNRKYTIKVEVPKGAWRSYLNMELTIPIFATNSDCELIVKAMQGLLKDGNPIPSAIAANSGIY*ANFTQFVLVDNGGTGDVTVAPSNFANGIAEWISSNSRSQAYKVTCS**VRRRG** |
| tdPCP | SKTIVLSVGEATRTLTEIQSTADRQIFEEKVGPLVGRLRLTASLRQNGAKTAYRVNLKLDQADVVDSGLPKVRYTQVWSHDVTIVANSTEASRKSLYDLTKSLVATSQVEDLVVNLVPLGRADPLA*SKTIVLSVGEATRTLTEIQSTADRQIFEEKVGPLVGRLRLTASLRQNGAKTAYRVNLKLDQADVVDSGLPKVRYTQVWSHDVTIVANSTEASRKSLYDLTKSLVATSQVEDLVVNLVPLR* |
| cpPCP-G48 | SKTAYRVNLKLDQADVVDSGLPKVRYTQVWSHDVTIVANSTEASRKSLYDLTKSLVATSQVEDLVVNLVPLGRADPLA*SKTIVLSVGEATRTLTEIQSTADRQIFEEKVGPLVGRLRLTASLRQNGAKTAYRVNLKLDQADVVDSGLPKVRYTQVWSHDVTIVANSTEASRKSLYDLTKSLVATSQVEDLVVNLVPLGR*AGALASKTIVLSVGEATRTLTEIQSTADRQIFEEKVGPLVGRLRLTAS**LRQNG** |

| | |
|---|---|
| sol-cpPCP-G48<br><br>Mutations in **yellow** | SKTAYRVNLKLDQADVVDSG**A**PKVRYTQVWSHDVTIVANSTEASRKSLYDLTKSLVATSQVEDLVVNLVPLGRADPLA*SKTIVLSVGEATRTLTEIQ**R**TADRQIFEEKVGP**A**VGRLRLTASLRQNGAKTAYRVNLKLDQADVVDSG**A**PKVRYTQVWSHDVTIVANSTEASRKSLYDLTKSLVATSQVEDLVVNLVPLG*RAGALASKTIVLSVGEATRTLTEIQ**R**TADRQIFEEKVGP**A**VGRLRLTAS**LRQNG** |
| dPCP-G48 | SKTAYRVNLKLDQADVVDSGAPKVRYTQVWSHDVTIVANSTEASRKSLYDLTKSLVATSQVEDLVVNLVPLGRADPLA*SKTIVLSVGEATRTLTEIQRTADRQIFEEKVGPAVGRLRLTASLRQNGAKTAYRVNLKLDQADVVDSGAPKVRYTQVWSHDVTIVANSTEASRKSLYDLTKSLVATSQVEDLVVNLVPLG*RAGALASKTIVLSVGEATRTLTEIQRTADRQIFEEKVGPAVGRLRLTAS**LRQNGRRRG** |
| dPCP-N47 | SKTAYRVNLKLDQADVVDSGAPKVRYTQVWSHDVTIVANSTEASRKSLYDLTKSLVATSQVEDLVVNLVPLGRADPLA*SKTIVLSVGEATRTLTEIQRTADRQIFEEKVGPAVGRLRLTASLRQNGAKTAYRVNLKLDQADVVDSGAPKVRYTQVWSHDVTIVANSTEASRKSLYDLTKSLVATSQVEDLVVNLVPLG*RAGALASKTIVLSVGEATRTLTEIQRTADRQIFEEKVGPAVGRLRLTAS**LRQNRRRG** |
| **dPCP-Q46**<br><br>**'dPCP'** | SKTAYRVNLKLDQADVVDSGAPKVRYTQVWSHDVTIVANSTEASRKSLYDLTKSLVATSQVEDLVVNLVPLGRADPLA*SKTIVLSVGEATRTLTEIQRTADRQIFEEKVGPAVGRLRLTASLRQNGAKTAYRVNLKLDQADVVDSGAPKVRYTQVWSHDVTIVANSTEASRKSLYDLTKSLVATSQVEDLVVNLVPLG*RAGALASKTIVLSVGEATRTLTEIQRTADRQIFEEKVGPAVGRLRLTAS**LRQRRRG** |
| dPCP-R45 | SKTAYRVNLKLDQADVVDSGAPKVRYTQVWSHDVTIVANSTEASRKSLYDLTKSLVATSQVEDLVVNLVPLGRADPLA*SKTIVLSVGEATRTLTEIQRTADRQIFEEKVGPAVGRLRLTASLRQNGAKTAYRVNLKLDQADVVDSGAPKVRYTQVWSHDVTIVANSTEASRKSLYDLTKSLVATSQVEDLVVNLVPLG*RAGALASKTIVLSVGEATRTLTEIQRTADRQIFEEKVGPAVGRLRLTAS**LRRRRG** |
| dPCP-L44 | SKTAYRVNLKLDQADVVDSGAPKVRYTQVWSHDVTIVANSTEASRKSLYDLTKSLVATSQVEDLVVNLVPLGRADPLA*SKTIVLSVGEATRTLTEIQRTADRQIFEEKVGPAVGRLRLTASLRQNGAKTAYRVNLKLDQADVVDSGAPKVRYTQVWSHDVTIVANSTEASRKSLYDLTKSLVATSQVEDLVVNLVPLG*RAGALASKTIVLSVGEATRTLTEIQRTADRQIFEEKVGPAVGRLRLTAS**LRRRG** |

*Each dimeric subunit is displayed as underlined and *italicized* text, respectively. Amino acids that link the subunits is shown in gray. Amino acids that serve as penultimate residues to degron fusion are **bolded**. Degron sequences are shown in **bold red** text. Mutations used to create the 'solubilized' cpPCP variant are highlighted in yellow.
